# sRNAfrag: A pipeline and suite of tools to analyze fragmentation in small RNA sequencing data

**DOI:** 10.1101/2023.08.19.553943

**Authors:** Ken Nakatsu, Mayumi Jijiwa, Vedbar Khadka, Masaki Nasu, Matthew Huo, Youping Deng

**Author notes:** Contributing authors.

## Abstract

Fragments derived from small RNAs such as small nucleolar RNAs hold biological relevance. However, they remain poorly understood, calling for more comprehensive methods for analysis. We developed sRNAfrag, a standardized workflow and set of scripts to quantify and analyze sRNA fragmentation of any biotype. In a benchmark, it is able to detect loci of mature microRNAs fragmented from precursors and, utilizing multi-mapping events, the conserved 5’ seed sequence of miRNAs which we believe may extraoplate to other small RNA fragments. The tool detected 1411 snoRNA fragment conservation events between 2/4 eukaryotic species, providing the opportunity to explore motifs and fragmentation patterns not only within species, but between. Availability: https://github.com/kenminsoo/sRNAfrag.

## 1 Introduction

Evidence of transfer RNA (tRNA) fragmentation has been reported since at least 1969 [1]. However, it was only recently that their functional roles in disease was revealed [2, 3]. Fragmentation of other small RNAs (sRNAs) such as small nucleolar RNAs (snoRNAs) and ribosomal RNAs (rRNAs) have also been reported, majority of which are derived from the 3’ and 5’ ends of the primary transcript [4]. While much of the sRNA fragment literature has been focused on tRNA derived fragments, there is rising evidence that fragments derived from other biotypes of sRNAs have a mechanistic role in disease and human health by influencing translation and alternative splicing [5–7]. Reanalysis of Photoactivatable-Ribonucleoside-Enhanced Cross linking and Immunoprecipitation (PAR-CLIP) sequencing data sets have revealed that fragments derived from snoRNAs and rRNAs associate with Argonaute proteins, a component of the RNA-induced silencing complex which facilitates RNA interference [8–10]. Furthermore, fragmentation of sRNAs has been found to occur in eukaryotes besides humans, including mice, fruit flies, and chickens [11]. Thus, a pipeline that can easily call fragments is needed to understand why fragmentation is conserved, which may provide insights into their function.

sRNA sequencing libraries do not undergo fragmentation (typical in standard RNA-seq) and are instead size selected through gel or bead based protocols. [12]. This is useful to keep in mind when designing computational pipelines and was used in Tang et. al. Anchor Alignment-based Small RNA Annotation (AASRA) method to justify the addition of anchors to the 3’ and 5’ ends of sRNA sequencing reads to discriminate between similar sRNAs [13]. With this in mind, we assume throughout this paper that if a portion of a longer sRNA (perhaps 100-200nt) has supporting reads, we count it as a fragment of that longer sRNA as many others have done [4–9].

One of the shortcomings in non-tRNA sRNA-fragment literature is that it does not have curated annotations such as MINTbase [14]. sRNAfrag seeks to be a step towards generating such annotations. MINTbase was generated using the MINTmap algorithm which creates a lookup table for every possible fragment that can be derived from tRNAs, assigning each to a unique sequence-based license plate allowing IDs to be reproduced easily [15]. Authors of the MINTmap algorithm recognized that generated fragments are sometimes found outside of the tRNA space, mapping to other locations in the genome, highlighting the importance of validating potential fragments before claiming their existence.

Other tools exist to analyze the fragmentation of sRNAs that are not tRNAs (note that our pipeline can analyze tRNA fragments). The FlaiMapper algorithm detects peaks within BAM files and can accurately reconstruct start and end loci for mature miRNAs [16]. However, since BAM files are used as an input, users must make decisions while processing raw reads which can lead to different results and introduced biases dependent upon factors such as the assignment of multi-mapping reads or mismatch parameters [17]. sRNAs including rRNAs, snRNAs, snoRNAs, and tRNAs also often have many isoforms in eukaryotic genomes that fragments align to [18–21]. Thus, the multi-mapping problem must at least be addressed.

The authors behind SURFr (Short Uncharacterized RNA Finder) present a new alignment algorithm and use continuous wavelet transformation to identify peaks [22]. While it represents a new and interesting way to handle fragments, their web portal could not be accessed as of June 16, 2023.

The AASRA method by Tang et. al. motivated us to apply size-selection knowledge to a pipeline [13]. The MINTmap algorithm provided a basis as to how fragments of longer sncRNAs could be annotated and brought to our attention the need to consider mappings to other places within the genome [23]. Finally, ideas from FlaiMapper and SURFr helped us to develop the intuition needed to accurately call start and end positions of fragments [16, 22]. We sought to combine elements from these different methods and pipelines to create sRNAfrag, a pipeline to analyze sRNA fragmentation not only from small RNA sequencing runs but also between them. The pipeline requires only raw reads, an annotation file, and a reference genome to run, leaving only demultiplexing of samples and library preparation to the user.

## 2 Results

### 2.1 sRNAfrag Effectively Calls Peaks and Counts

sRNAfrag extracts both counts and potential start/end loci for users. Acting as a ground truth for fragmentation, mature miRNA loci (start and end positions) were extracted from MiRbase. We considered pre-miRNAs as the source transcript in this benchmark. sRNAfrag had 597 pre-miRNAs with supporting fragments after filtering compared to the 642 that FlaiMapper obtained. The discrepancy between the two pipelines exist because sRNAfrag deals with multi-mapping by dividing the number of associated counts by the number of loci it could have originated from. Thus, filtering based on counts is more strict in sRNAfrag. Of these pre-miRNAs, sRNAfrag called 94.7% and 86.5% of start and end loci within 1 bp of their true position (Fig 1A). This is comparable to FlaiMapper which called 92.4% and 85.2% of start and end loci in our dataset with default settings [16]. Notably, offsets are not a result of false peak callings. Rather, count peaks exist at positions that do not exactly match loci in miRbase as can be seen in [Additional Figure 1].

**Fig. 1.**
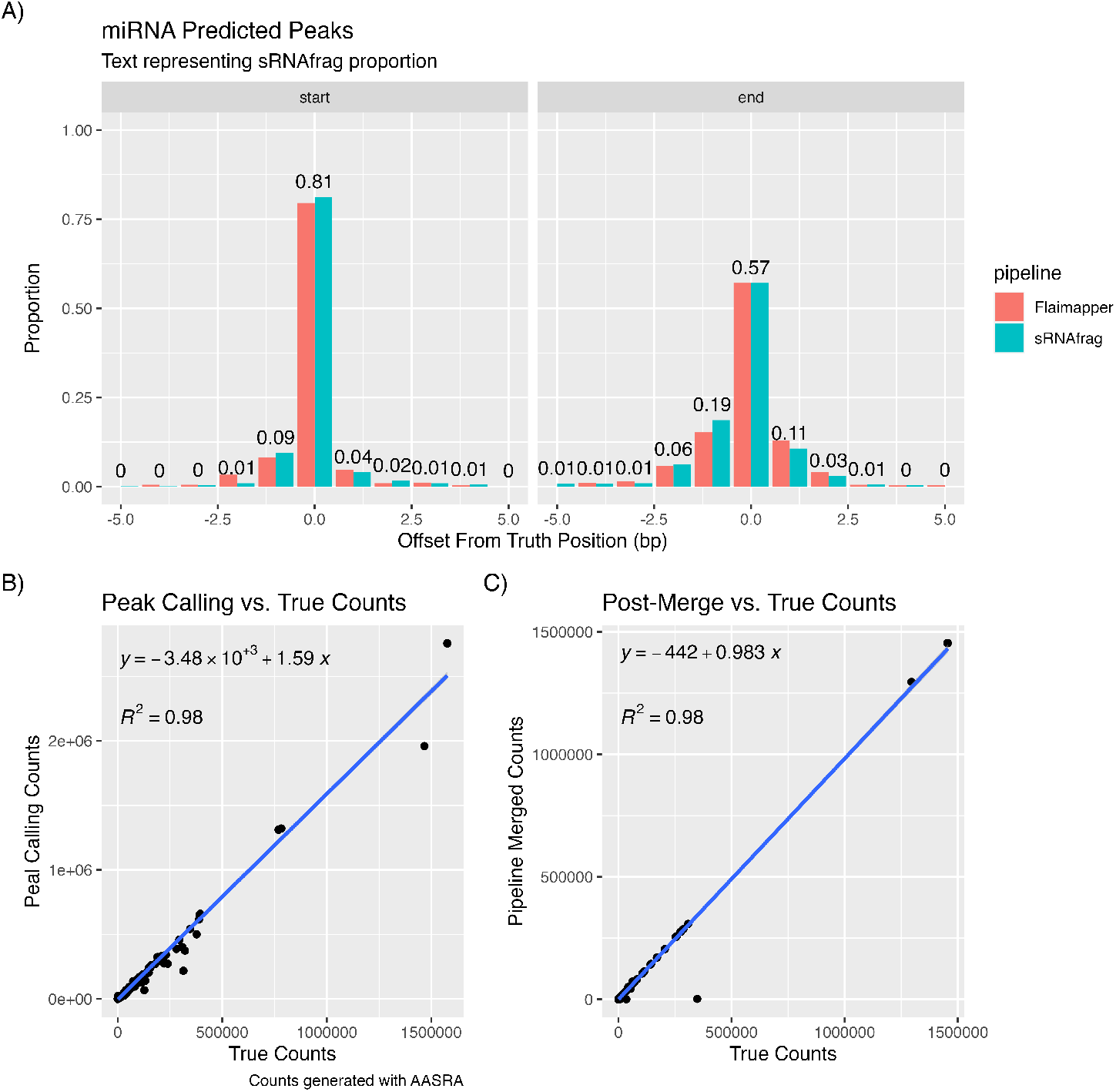
Peak and Count Calling Performance. A) Proportion of detected peaks vs. Offset from truth for end and start loci. B) Peak calling counts vs. AASRA generated counts. Counts are unmerged and represent the unadjusted counts for that loci. C) Merged counts after P2 Module vs. AASRA generated counts.

Comparing the counts used to call peaks to that of AASRA shows that sRNAfrag utilizes counts that are comparable to more standard small RNA-seq workflows during peak calling, affirming that, in our pipeline, loci-count relationships are properly preserved and used (Fig 1B). A benchmark of the AASRA method was conducted on a synthetic dataset presented in [Additional Figure 2]. However, using these raw counts for downstream purposes would produce biased results. Thus, we show that when counts are merged (fragments that are clustered together), results are closer to that of AASRA (Fig 1C). We infer that the underestimation of slope (0.983) and negative intercept is partially due to the AASRA pipeline’s behavior to assign alignments that have a 1 n.t. mismatch stretching beyond the source transcript due to the anchoring behavior. Because of this, starting positions prior to the first loci of the annotation are counted as being apart of that transcript whereas sRNAfrag does not as a sequence preceding a transcript is not considered when generating the lookup table.

### 2.2 Multi-Mapping Events Reveal Potentially Important Conserved Fragments Loci

The utility of sRNAfrag expands beyond the basic analyses presented in the summary report depicted in [Additional File 1].In the small RNA space, the multi-mapping problem poses a formidable problem. However, we decided to utilize multi-mapping reads to attempt to elucidate important loci. We again explore miRNAs as a model. The 5’ seed sequence of miRNAs represent the functionally important portion of the molecule [24]. Furthermore, miRNA families, arising from duplications events, likely are subject to evolutionary pressures to retain the same seed sequence [25]. Utilizing a function packaged with sRNAfrag, we extracted the a) most commonly shared fragment(s) between source transcripts and b) most counted fragment within a cluster. The mir-30 family had four (3 are depicted, with the final being 1 n.t. longer) fragments that were shared amongst all potential sources (Fig 2A). Each were aligned at their 5’ ends, representing the beginning of the 5’ derived mature miRNA (hsamiR-30c-5p). We investigate all clusters of fragments with more than one potential source transcript and found that the 5’ end of the most counted fragment match the most commonly shared fragment 65.9% of the time (Fig 2B).

**Fig. 2.**
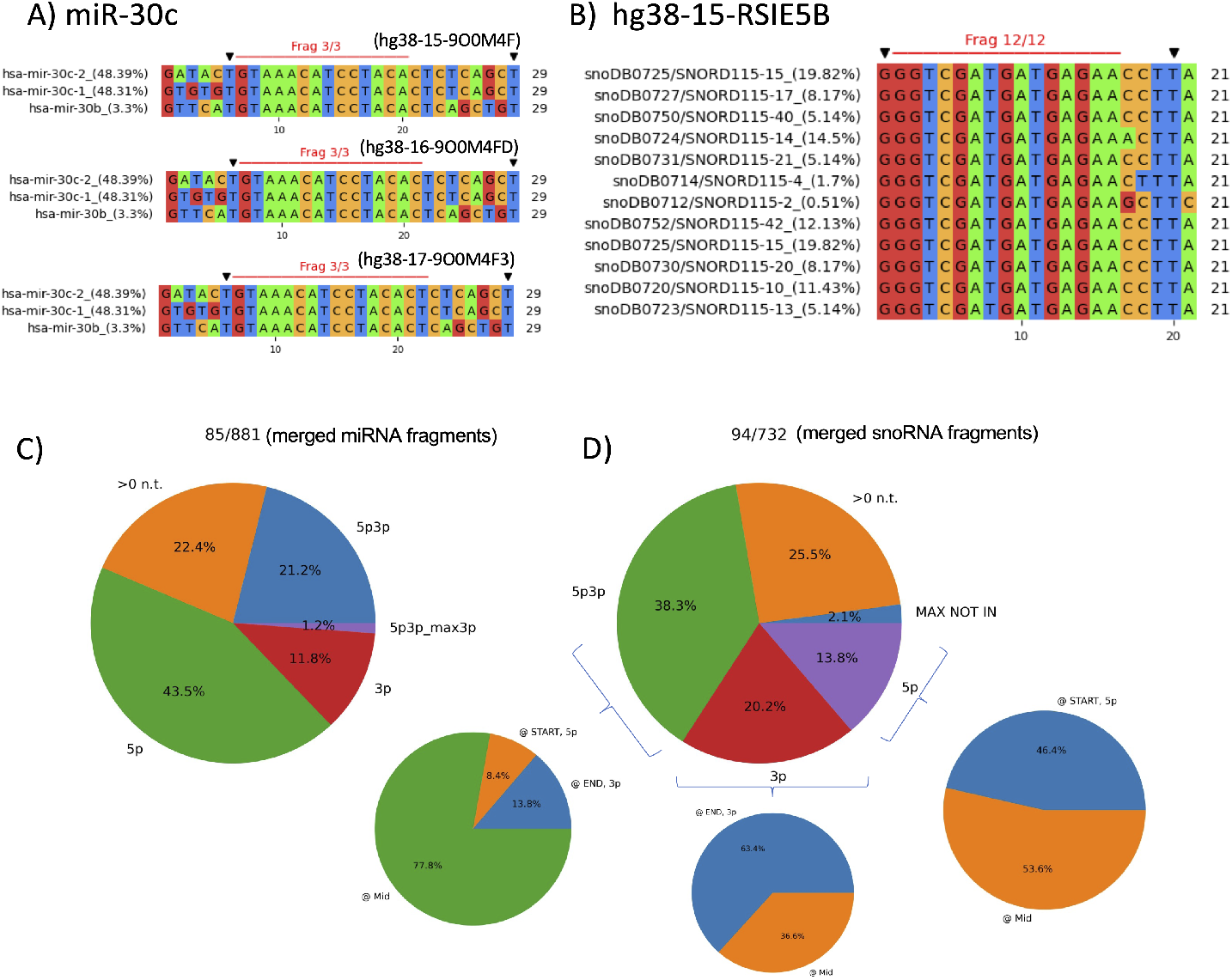
Comparative Analysis of Fragments Involved in Multi-Mapping Events. A-B) Black triangles depict the most highly expressed fragment of the source transcript. The red line depicts the most shared fragment between potential sources. Three clustered fragments shared between has-miR-30b/30c and one fragment shared between 12 isoforms of snoRD115 whose 5’ end is the same as the most conserved fragment is depicted. C-D) Piechart depicting the proportion of merged fragments (with more than one possible source transcript) whose most commonly shared fragment between sources has a the same 3’ or 5’ loci as the most counted transcript. The percentage of transcripts whose shared loci represents either the start or end of the source transcript is also indicated. Greater than 0 represents occurrences when the most found and counted transcript do not match.

We attempted to apply the same logic to a fragment derived from snoRD115. The fragment in question was shared between 12 source transcripts that had slight differences in sequence (Fig 2B). While the proposed MSA appeared promising, we found that the fragmentation of snoRNAs was inherently different from miRNAs. Exploring the location of these fragmentation events showed that 40% of conservation events were at terminal portion of the source transcript compared to just 1.5% for miRNAs (Fig 2D). In the future, similar assignments could allow for better understanding of fragment biogenesis in other biotypes of small RNAs.

### 2.3 License Plating Allows Conserved Fragments to be Detected

Given that orthologous snoRNA genes are fairly common, we decided to investigate if fragments of similar sequence could also be conserved across species [26]. We have written and documented command line functions in our public repository to detect similar fragments between runs, utilizing hamming distance as a measure of similarity. Hamming distance calculates the number of differences between two sequences of equal length. Using this method, 1411 conservation events between two species were found with a hamming distance at most of 3, majority of which were fragments shared between M. musculus and H. sapiens (Fig 3A).

**Fig. 3.**
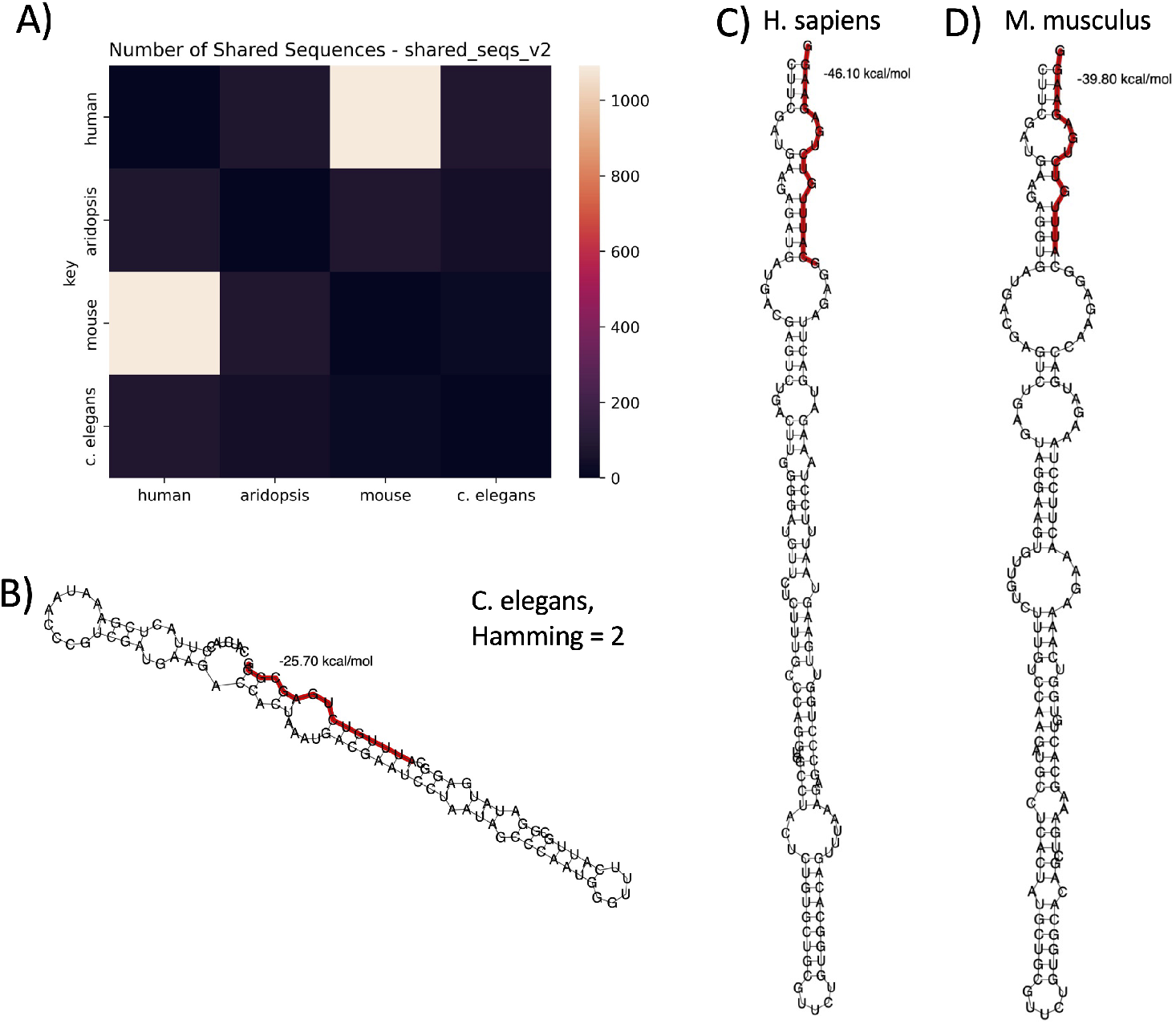
Shared snoRNA Fragments between Species. Source transcripts are depicted with fragment source cluster loci highlighted in red. **A)** Heatmap depicting number of shared fragments between four species, M. musculus (mm), H. sapiens (hs), A. thaliana (at), and C. elegans (ce). Diagonals are set to 0. **B)** Shared snoRNA fragment with hamming distance of two relative to H. sapien and M. musculus fragments in panel (C) and (D). Derived from unassigned transcript 14429. **C)** H. sapiens snoRNA derived fragment, with hamming distance of 0 relative to M. musculus fragment depicted in panel (D). Derived from SNORD15A. **D)** M. musculus snoRNA derived fragment. Derived from NR 002172.1 (SNORD15A).

Out of the 1411, we chose to investigate one shared fragment derived from the 3’ ends of primary transcripts in 3/4 of our chosen species. RNAfold and RNAplot was used to generate minimum free energy structures of source transcripts [27]. The sRNAfrag workflow associates fragments with a cluster which has associated start and end loci. The cluster start and end are highlighted on the secondary structure, as opposed to the fragment start and end (Fig 3B-D). The fragment derived from the C. elegans snoRNA was the most different out of the three, with a hamming distance of two (Fig 3B). The fragments derived from snoRNAs in H. sapiens and M. musculus shared the same sequence (Fig 3C-D). Fragments were derived from SNORD15A which is annotated for M. musculus and H. sapiens. This is a proof of concept that attempting to explore fragmentation between species may not only be fruitful, but is also made easier with sRNAfrag as its outputs can be joined and used in scripts such as the one used in this section.

### 2.4 sRNAfrag Run Time Scales Linearly for Multiple Biotypes and Species

We tested sRNAfrag with four different species (M. musculus, A. thaliana, C. elegans, H. sapiens) and three biotypes of sRNAs (snoRNA, snRNA, rRNA) with a maximum of one job. The amount of time that sRNAfrag takes exhibits a linear relationship with the number of fragmnts discovered in a manner that is mostly independent of the number of sequenced reads (Fig 4A). For every 1000 additional unique fragments discovered, users can expect an additional 23 seconds of run time. Note that parallelization can increase speed of time intensive modules such as sorting of SAM file and alignment back to reference genome. This is slower than reported times in SURFr, taking an average of 4.5 minutes to complete jobs [22]. However, our pipeline realigns fragments back to the reference genome which takes significant amount of time.

**Fig. 4.**
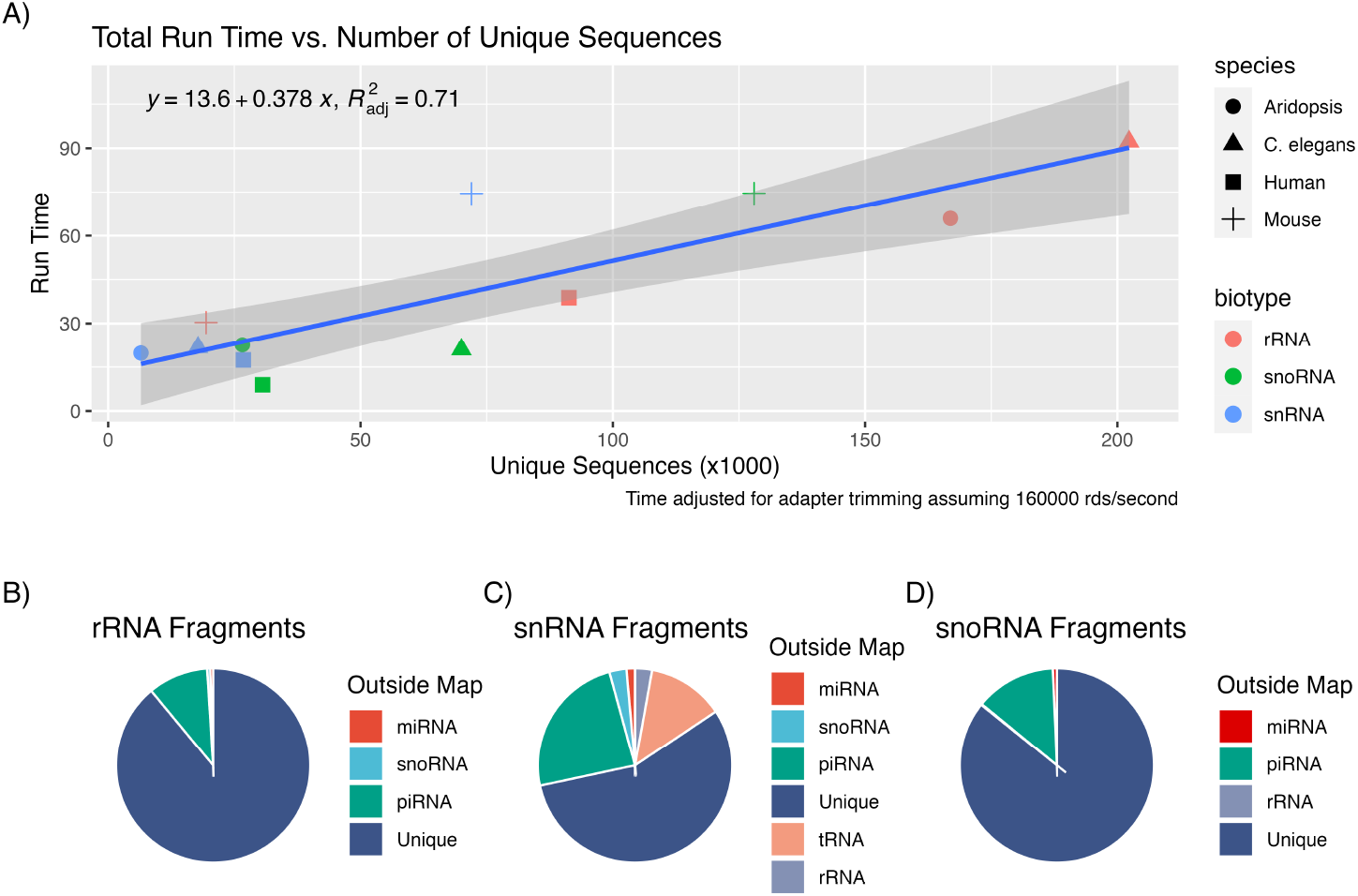
sRNAfrag Run Time and Human Outside Map Composition. A) Overall run time in minutes. B-D) The composition of out of space maps for rRNA, snRNA, and snoRNA fragments in humans.

Especially large run times were observed for rRNA fragments, which is unsurprising given that the pipeline processed 20,000-200,000 fragments. This is because when the number of fragments surpasses 8000, sampling of the smallest fragments is conducted, which is slow. Given that discovering outside maps take time, we investigated where fragments are mapping to in our cell line dataset. We found that most reads from fragments map to piRNAs (Fig 4B-D). Interesting, snRNA U1 has been found to be similar to that of tandem repeat tRNA which may explain why snRNA fragment alignments were annotated as tRNAs [28]. We believe that these results indicate that outside maps should always be assessed as they are always existent.

Generation of the lookup table takes time as it requires that sequences be sorted and queried. We originally generated a lookup table of all possible fragments. However, this led to memory issues, especially as possible fragments would range in the tens of millions (Table 1). Furthermore, fragments that exist typically represent only a small fraction of possible sequences in nearly every species tested.

**Table 1.** Possible maximum number of fragments generated by provided annotation files and percentage found in test data.

| Species | Biotype | Possible Fragments (thousands) | % Detected |
| --- | --- | --- | --- |
| H. sapiens | rRNA | 5827 | 2% |
| H. sapiens | snoRNA | 5762 | 0.5% |
| H. sapiens | snRNA | 47338 | 0.06% |
| A. thaliana | rRNA | 544 | 31% |
| A. thaliana | snoRNA | 573 | 5% |
| A. thaliana | snRNA | 350 | 2% |
| C. elegans | rRNA | 252 | 80% |
| C. elegans | snoRNA | 1004 | 7% |
| C. elegans | snRNA | 316 | 6% |
| M. musculus | rRNA | 224 | 9% |
| M. musculus | snoRNA | 3067 | 4% |
| M. musculus | snRNA | 2204 | 4% |

**Table 2.** Entries in the lookup table contain information regarding source transcript and the position from which it originates from. The start locations are standardized to fall between 0-1 and averaged if more than one position exists. Additional columns not shown contain information about 5’ and 3’ flanking sequences. If a fragment can originate from more than one source in the provided annotation, all information is delimited with a special character in the sources column.

| seq | sources | num | avg start | std | ID |
| --- | --- | --- | --- | --- | --- |
| ACG... | 3;25_S1<sup>1</sup>_-...0.008 5;27_S2_-...0.02 | 2 | 0.012 | 0.004 | prefix-23-E19OW6PI0P |

<sup>1</sup>A source, i.e. SNORD3A.

The upper maximum number of fragments that could be generated in the lookup table can be calculated with equation 1 when *m* is maximum length, *k* is minimum length, *i* is the ith source transcript, *j* is the number of source transcripts, and *n* is the length of the transcript:

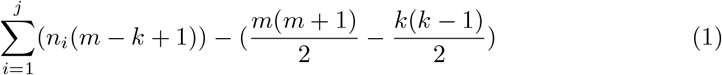

Intron-derived fragments that are outside of the 20-22 bp range of miRNAs may be interesting to investigate as fragments of different sizes have been found to be potential sources of biomarkers for disease [29]. Lopes et. al. reported that transcript length peaks at 2,065 bp over 92,696 transcripts and genes [30]. Analyzing fragmentation over 20000 protein coding genes would yield a theoretical number of 1.3 billion possible fragments. Assuming that fragmentation of protein-coding gene follows the trend of snRNAs (0.06% of fragments out of all possible) for a total of 780,000 detected fragments, this would take about 5 hours to run. We previously found that lookup table generation occurs at a rate of 11,000 fragments / second. Ignoring memory limitations, lookup table generation alone would take 32 hours and would take up 252 GB of space. Thus, our approach of sorting sequences for lookup table creation allows for wider scalability and application.

## 3 Discussion

The development and validation of sRNAfrag presents an opportunity to advance our understanding of small RNA fragmentation by providing a framework to explore multi-mapping events and conserved fragments between species. It packages together tools to generate figures that are relevant to small RNA fragmentation beyond simple analyses. Here, we will discuss why we did not choose to have sRNAfrag generate annotations, limitations, and future directions for the tool.

### 3.1 Generation of Annotations?

Unlike sRNAfrag, Flaimapper generates annotations for users, which is admittedly convenient. However, we believe this is dangerous for three primary reasons:

1. Calling start and end loci solely on peaks is subject to its limitations as can be seen by the similar performance between FlaiMapper and sRNAfrag (Fig 1A).
2. miRNA sequences derived from precursors with both sufficient coverage and the proper secondary structure are predicted to have a false positive rate of 65% [31]. Similar issues likely exist with the discovery of small RNA fragments.
3. We should collaborate to develop annotations for fragments instead of allowing researchers to develop annotations independently. Currently, the tags “small RNA,” “snoRNA,” “snRNA,” “rRNA,” and “smRNA” yield only a total of 13 databases compared to 107 for “miRNA” in the National Genomics Data Center Database Commons [32].

Given these constraints, one may wonder, “what is the point of sRNAfrag?” On the surface, it certainly does what it proclaims to do, facilitate the analysis of small RNA fragmentation. See [Additional File 2] for how base sRNAfrag outputs can corroborate past findings by investigators such as Taft et. al. and Li. et. al [4, 11]. At its core, however, sRNAfrag seeks to be a framework that engages scientists interested in the Evolutionary, Structural (RNA), Algorithmic, Graphical, and Systems components (and more) of bioinformatics to attract a high impact community [33]. We believe such a pursuit possible because of novel discoveries continuing to occur in the non-tRNA fragmentation space with dates ranging from 2009-2022 [6, 7, 9, 11, 16, 34, 35]. While ambitious and subjective, this represents the core reason why sRNAfrag does not output annotations.

### 3.2 Limitations

The merging of fragments into clusters makes it difficult to ascertain which standard normalization methods to use. To remedy this, we have integrated three normalization methods, ratio normalization with miRNAs, transcripts per million, and counts per million (reads) [36]. This is discussed in further detail in [Additional File 3].

While this offers a method to normalize between samples in one run of sRNAfrag, it does not offer a comprehensive method to compare counts between runs. Combining simply on license plates is possible; however, we were wary to implement such an option as fragmentation analysis is likely subject to large technical variations based upon runs [37]. When preparing libraries for Illumina sequencing, size distributions can vary between size selection protocols such as gel electrophoresis, AMPure XP, Caliper Labchip XT, or Sage Science Pippin Prep [38]. A library that has a larger composition of shorter reads of lower complexity will lead to lower resolution when clustering fragments. Thus, in this first release of sRNAfrag, we only provide tools to qualitatively compare fragments between runs.

Finally, our peak calling algorithm does not account for secondary structure or motifs that may be indicative of true fragmentation sites. Thus, users are left with very little information as to whether or not their detected fragments are degradation products or not. Including motifs may make our pipeline more effective in detecting true fragments. For example, C/D box snoRNA derived fragments often contain their respective motifs (C: UGAUGA or TGATGA in the figure), as seen in Figure 2B. The C/D box motif has been reported to direct 5’ capping, thereby increasing the stability of snoRNAs which may explain why the C/D box is found fragmented at higher rates from the 5’ end of the parent transcript [11, 39].

### 3.3 Future Directions

Besides optimization and bug fixes, future development of sRNAfrag will focus on the detection of post-transcriptional modification events. In eukaryotic organisms, RNAs are degraded through 3’ exonuclease activity by the exosome complex (not to be confused with extracellular vesicles) [40]. We hypothesize that protected fragments hold greater biological significance. The 3’ end of hsa-miR-122 is subjected to a non-template addition of an adenine that can increase its stability [41]. We are curious if a similar mechanism exists for small RNA derived fragments. By pursuing this, we hope to unveil a larger number of potential fragments that may be significant to life and biology.

## 4 Conclusion

The sRNAfrag pipeline comprehensively and quickly executes to describe sRNA fragmentation in small RNA-seq data. The outputs are easy to understand and to work with. Counts are tabularly summarized and potential fragment sequences are outputted in a fasta format. Because of the variety of outputs, the sRNAfrag pipeline can be utilized beyond standard differential expression measures (which could be accomplished with packages such as DESeq2) [42]. For example, by mining cross-linking and immunoprecipitation (CLIP) sequencing datasets, one can identify what protein complexes are formed by these fragmented RNAs [43]. One could imagine Fig 6A, with one standard small RNA-seq dataset and three CLIP-seq datasets instead of species to detect which fragments associate certain proteins. Knockdown experiments can also be used to elucidate potential mechanisms of their biogenesis [44].

There are also many tools that can predict function of small RNAs based on their sequence [45]. sRNAfrag outputs can easily be modified with scripts to fit inputs for other methods. For example, a user could recursively loop through two fasta files, one with mRNAs and the other with outputs from this pipeline to generate the proper input for TargetNet, a deep learning target prediction method [46].

We believe that sRNAfrag will be helpful in elucidating not only potential biomarkers for disease, but also for investigating the role of RNA fragmentation in larger contexts. Eventually, we would like to use sRNAfrag to develop a standardized method to curate a database of high-confidence sRNA fragments. Such developments are out of the scope of this paper and will be addressed in future versions of sRNAfrag as we strongly believe that inaccurate annotations are worse than none.

## 5 Methods

### 5.1 Pipeline Overview and Annotation Data Sources

sRNAfrag can be broken down into three distinct parts which we have labeled as P1, S1, and P2 (Fig 5). Users specify the location of files, adapter sequences, which modules to run, etc. by modifying a YAML configuration file. YAML files are similar to JSON files. A modified annotation file in General Transfer Format (GTF), along with a folder of gzipped fastq files is used as an input.

**Fig. 5.**
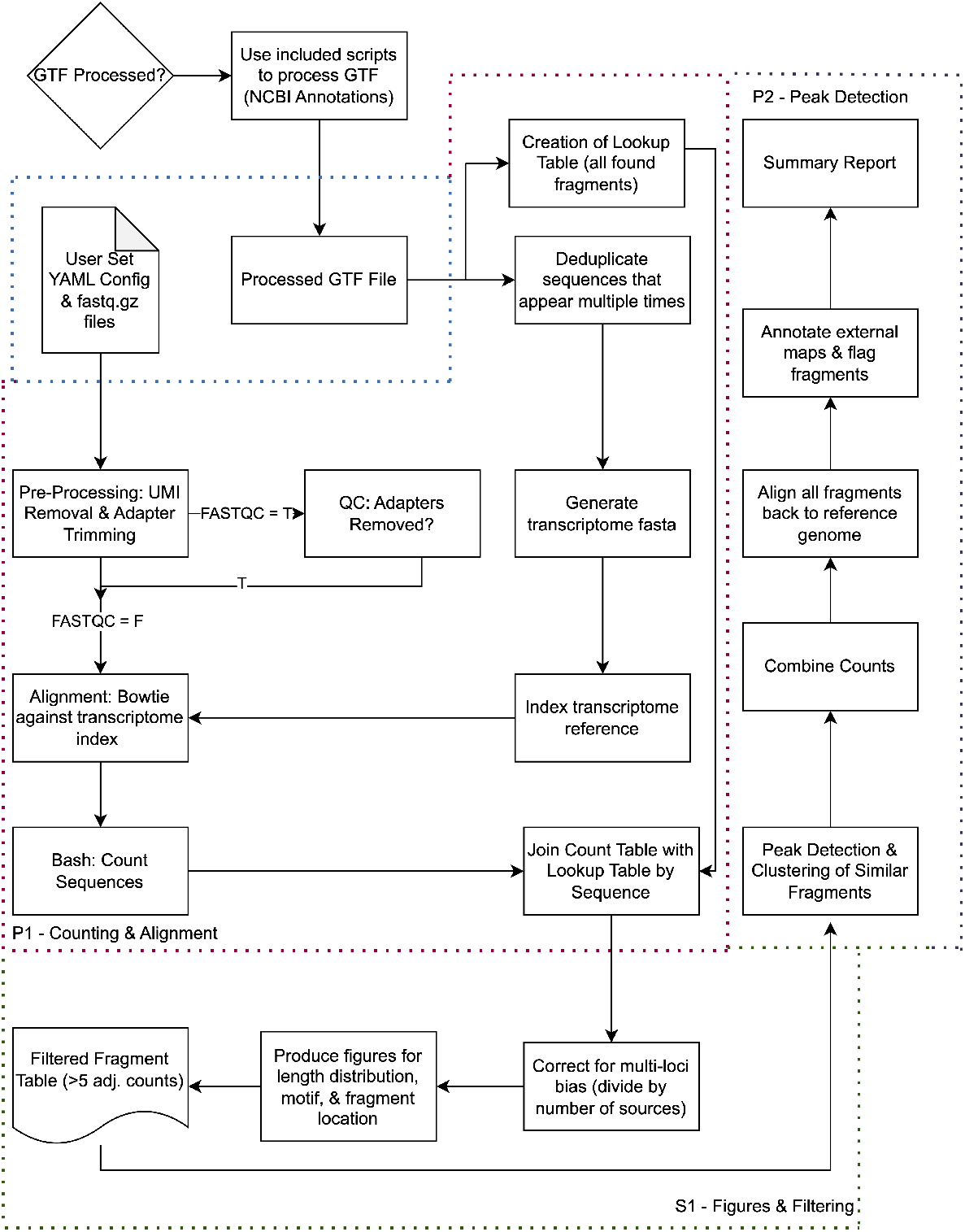
Pipeline Workflow. Summary of the sRNAfrag pipeline. Users first configure the pipeline by modifying an included YAML file. Users may also modify annotation files using included scripts if needed. Three modules, P1, S1, and P2, are used to processed raw gzipped fastq files to a final summary report while also generating figures.

We have created and provided scripts that parse NCBI annotations. This was applied to three common model species, M. musculus, C. elegans, and A. thaliana [47]. Human annotations were derived from snoDB (snoRNA), RNAcentral (snRNA), and Integrated Transcript Annotation for small RNA (rRNA) [48–50]. All processing is documented on our public repository (https://github.com/kenminsoo/sRNAfrag). Methods for validating and combining annotations from multiple sources are discussed further in [Additional File 4] which can improve sRNAfrag’s utility. GNU Parallel accommodates for parallel processing [51].

### 5.2 P1 Module -Initial Processing

Raw reads are processed for UMIs and adapters with UMI tools and AdapterRemoval [52, 53]. A subset of 2 Mb of read data is passed into FastQC to ensure that adapters were removed. Reads are then aligned to a transcriptome with bowtie using the following parameters [54]:

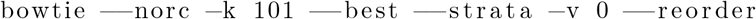

Aligned reads are extracted using SAMTools and sorted using unix commands [55]. Next, a lookup table of found fragments is generated using the modified GTF file. Pre-processing of the annotation adds sequence information with BEDTools which is needed for the generation of the lookup table [56]. An entry in the lookup table is shown in table 1.

License plate IDs are generated based on sequence, allowing for lookup tables generated from different annotations to share the same IDs, using code from the makers of MINTmap [15]. Then, the lookup table and each sample’s counts are joined on the basis of sequence.

### 5.3 S1 Module -Multi-Mapping Correction & Basic Analyses

An R script parses counts, dividing them by the number of potential fragment sources to account for multi-mapping. A linear regression is applied to the counts before and after the correction to allow users to assess its performance. A line graph depicting count density by length is also generated. ggseqlogo generates logos for each fragment length to detect potential motifs [57]. Transcripts with under 5 adjusted counts are filtered out prior to downstream analysis. Corrected and raw counts are saved to the specified out directory.

### 5.4 P2 Module -Merging & Compiling Fragments

Bipartite star graphs are generated to construct count peaks by loci on possible source transcripts with the python package NetworkX [58]. Formally, graphs are built with center nodes from S, the set of possible sources, and edges extend to nodes from F, the set of fragments, if its sequence can be found within a particular source node or transcript. Each edge holds count and loci data.

Conceptually, this is a standard peak calling problem. Thus, maxima are detected upon a positive to negative sign change. Smoothing is applied to these differences and can be defined formally as follows with source length *k*, loci *p*, and counts *n. p* = 0 is assumed to have 0 counts. A pandorable (the Pandas package) implementation of the algorithm is discussed in detail in [Additional File 5].

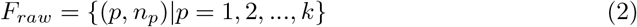

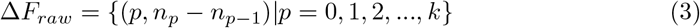

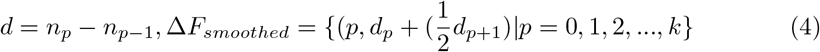

Equation 2 defines the distribution of counts by loci. Equation 3 and 4 represent the rate of change and smoothed rate of change, respectively. The smoothed rate of changed reduces the sensitivity of peak calling, as can be observed at loci 7 in Fig 6A. Sandwiched zeros are eliminated by assigning it to the count of the next loci for peak calling purposes and is visualized at loci 6 in Fig 6B.

**Fig. 6.**
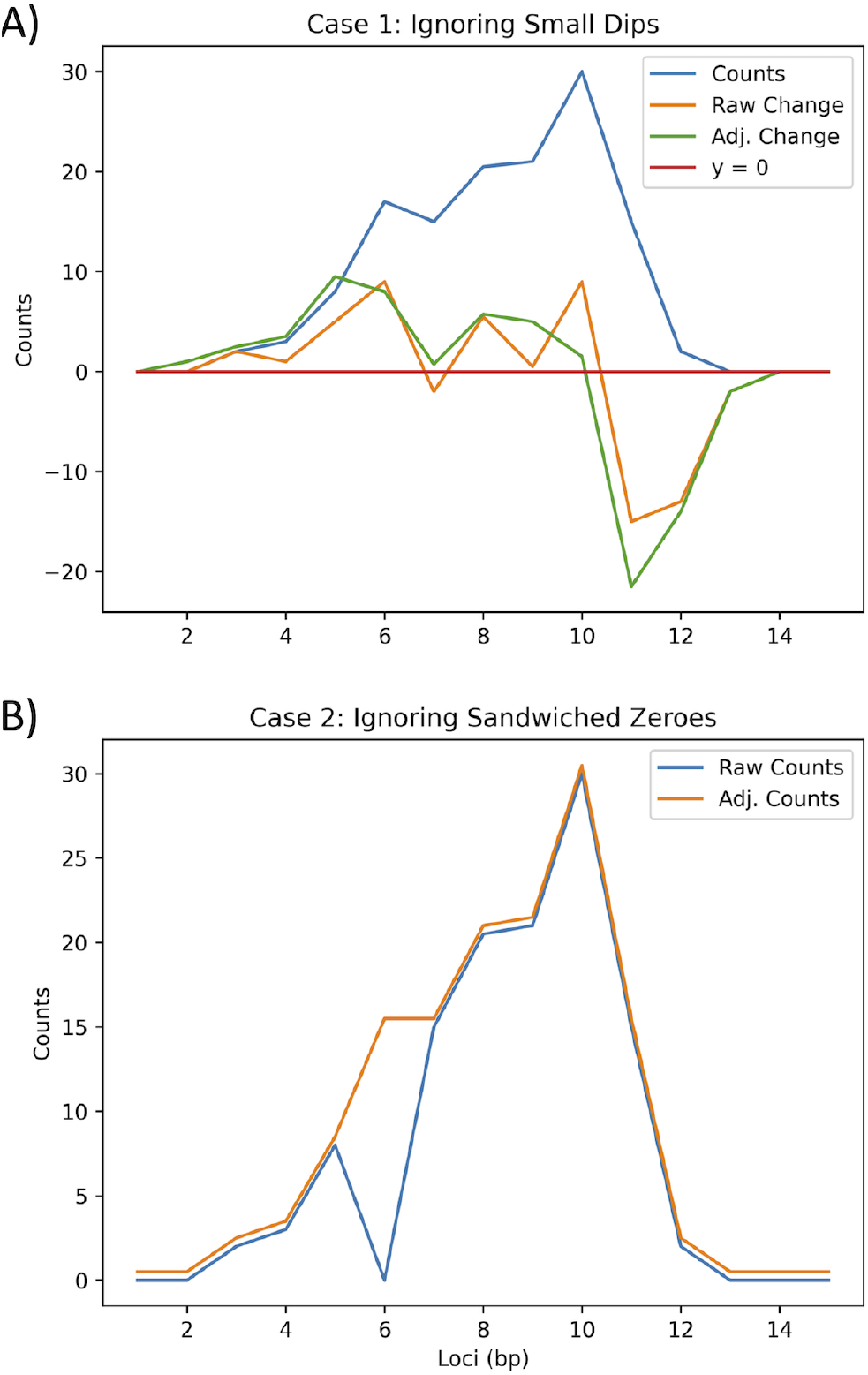
Addressing The Smoothing and Sandwich Problem. A) Depiction of the adjustment with the smoothing function which prevents peaks from being called if large peaks exist close by. B) Sandwiched zeros are assigned to the same number of counts as the next loci under the assumption that sandwiched zeros tend to be a result of sequencing noise.

Start and end peaks are called using the smoothed rate of change whenever a positive to negative sign change is detected. Then, for each source transcript (*S*_*n*_), fragments are clustered with minimum length of 10 (Fig 7).

**Fig. 7.**
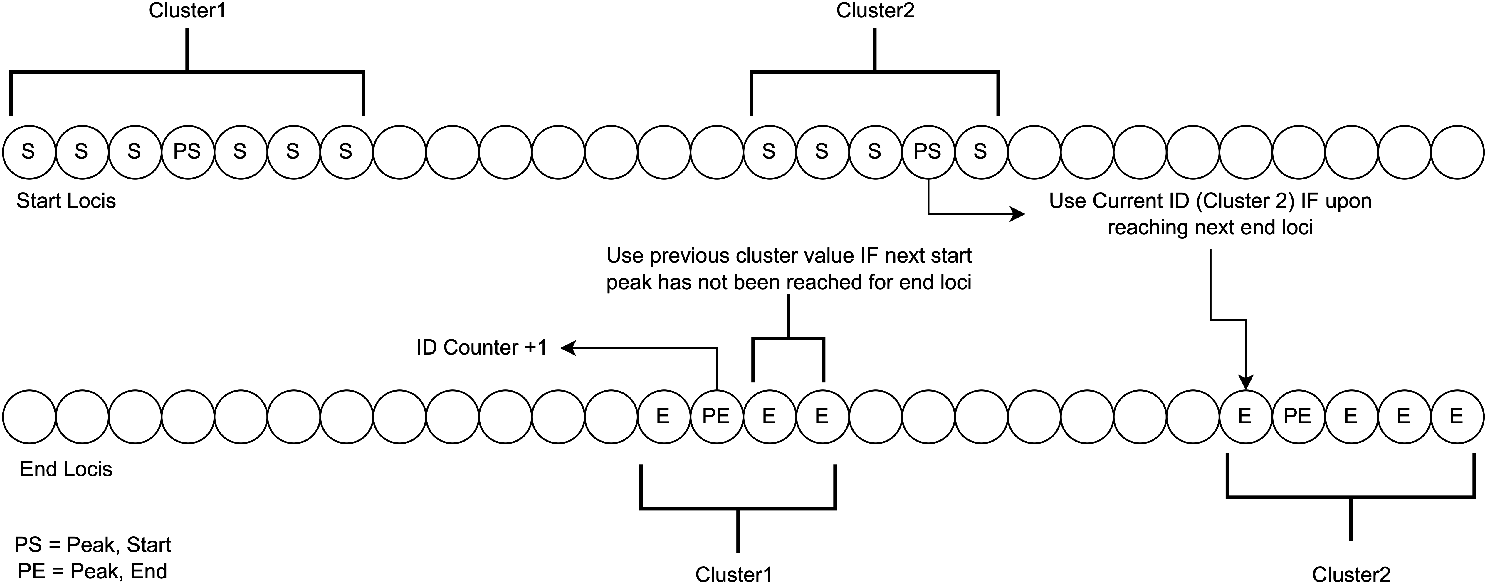
Clustering Algorithm. An example transcript is depicted with start and end loci labeled onto the sequence with detected peaks. Clusters are assigned using logic. Clusters must be at least 10 n.t. long before the id counter increases.

Cluster IDs are assigned to each loci. Fragment IDs (license plates) that associate with each source are assigned to a particular cluster. A clustering graph is initialized with fragment IDs as nodes. Edges are drawn if, on a particular source, a fragment shares the same start and end cluster ID. Independent subgraphs are extracted and all fragment counts are combined.

All fragments are aligned back to a reference genome to detect false positives. Unix commands are used to parse the alignments. FeatureCounts is used to annotate each fragment [59]. If there are over 8,000 fragments, fragments are randomly sampled by cluster to allow featureCounts to execute properly.

### 5.5 Summary & Outputs

A summary output converts a markdown file to an HTML summary report. Summary reports include all primary figures, allowing for quick interpretation of the data. Structure of the final directory is shown in Code 1.

#### Code 1

Pipeline Output

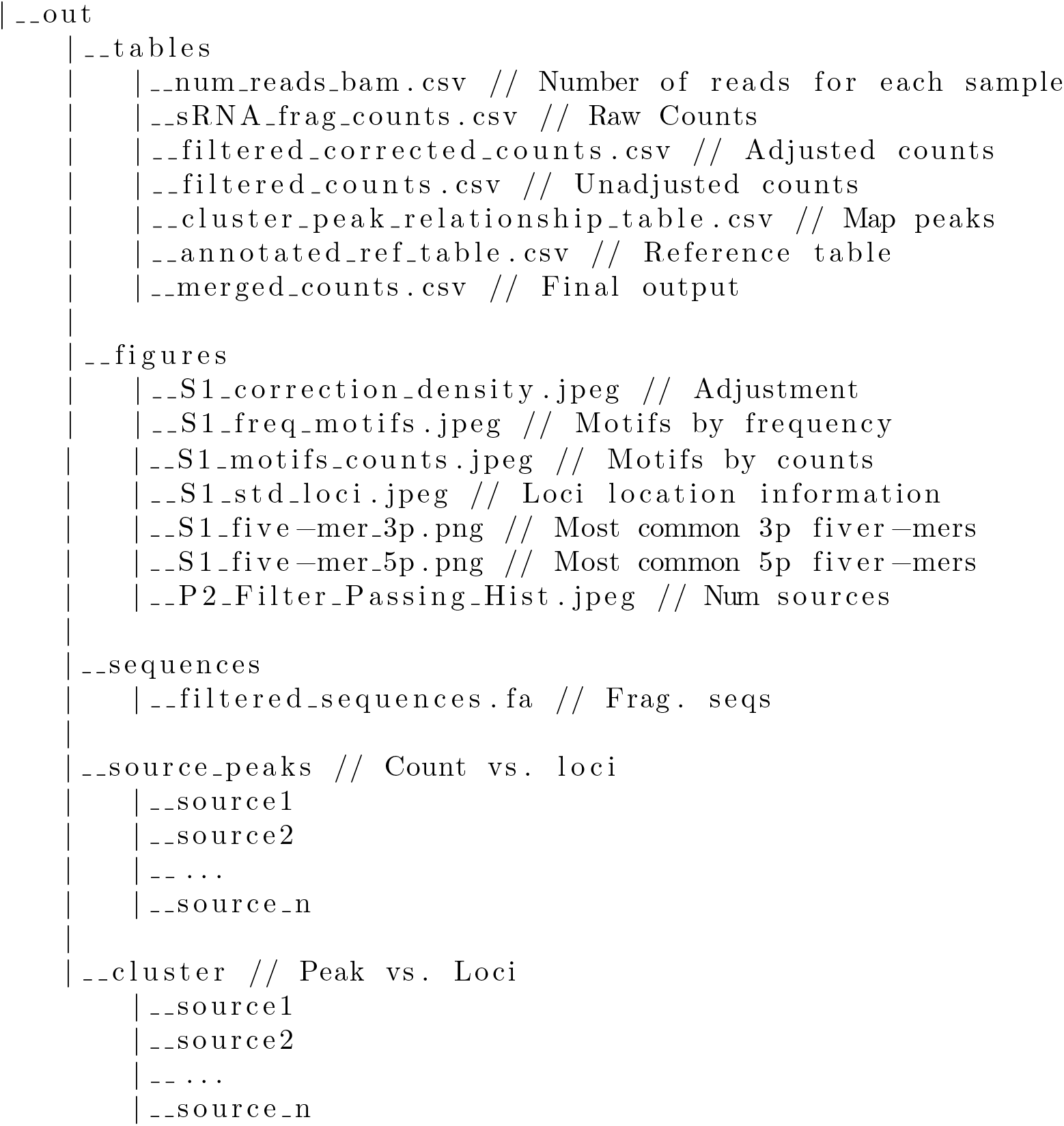

### 5.6 Pipeline Demonstration and Benchmarking

Public datasets for small RNA sequencing read for M. musculus (GSE213097 [60]), C. elegans (GSE192793 [61]), and A. thaliana (GSE152909 [62]) were used to demonstrate pipeline utility across species. We sequenced nine human cell line samples to demonstrate miRNA-loci calling. Sample information is included in table 3.

**Table 3.** Sample Information.

| Source | Sample Type | Notes | Species | n |
| --- | --- | --- | --- | --- |
| This Study | Cell Line | HMEC | *H. sapiens* | 1 |
| This Study | Cell Line | SK-N-SH | *H. sapiens* | 2 |
| This Study | Cell Line | IMR32 | *H. sapiens* | 2 |
| This Study | Cell Line | T47D | *H. sapiens* | 2 |
| This Study | Cell Line | HCC1143 | *H. sapiens* | 2 |
| GSE152909 | Seedling | amiR-SUL / ab57 | *A. thaliana* | 6 |
| GSE192794 | Whole animal | RFK1256 / N2 | *C. elegans* | 6 |
| GSE213097 | Plasmacytoid dendritic cells | IMQ Induced / Control wt | *M. musculus* | 6 |

To assess the peak calling algorithm, an miRNA benchmark was conducted using data generated from this study using miRbase as a ground truth [63]. Count accuracy for our pipeline was obtained with the AASRA pipeline [13].

Multi-species and biotype run-time metrics were obtained by running the pipeline with different genomes and annotations on a 2021 Macbook Pro, Apple M1 Max, with one core and 52GB of memory. For identifying conserved fragments across multiple species and plotting sRNA fragment source region secondary structure, command line functions apart of the sRNAfrag pipeline were used. RNAfold from the ViennaRNA package was used to generate the Minimum free energy structure [27]. PyMSAviz was used to visualize sequence alignments (https://pypi.org/project/pymsaviz/). All visualizations (except for time metrics) can be generated using scripts and configuration files in the “scripts” folder of our repository.

### 5.7 Cell Culture

SK-N-SH, IMR32, T47D, HCC1143, and HMEC cells were purchased from the American Type Culture Collection and maintained at 37°C in a humidified atmosphere of 5% CO2. SK-N-SH and IMR32 cells were grown in Dulbecco’s modified Eagle’s medium supplemented with 10% fetal bovine serum and 1% Penicillin-Streptomycin. T47D and HCC1143 cells were grown in ATCC-formulated RPMI-1640 Medium (ATCC) supplemented with 10% fetal bovine serum and 1% Penicillin-Streptomycin, and 0.2 Units/ml of human recombinant insulin (Life Technologies) for T47D. To maintain HMEC, Mammary Epithelial Cell Basal Medium (ATCC) supplemented with Mammary Epithelial Cell Growth Kit (ATCC) was used.

### 5.8 Total RNA Isolation from Cells

Total RNA was isolated from cells using miRNeasy Mini Kit (QIAGEN). Briefly, cells grown at 80% confluence in a 100 mm cell culture dish were washed with PBS and collected in a 1.5 ml centrifuge tube. Cells suspended in approximately 200ul of PBS were sonicated under RNase-free conditions. Then, the lysate was treated with 700 ul of QIAzol (QIAGEN) and 140 ul of chloroform to isolate nucleic acids in the aqueous phase. RNA was extracted following the manufacturer’s instruction and eluted with 30 uL of nucleic acid-free water. Finally, one ul of RNase inhibitor was added to the solution. The RNA quality and concentration were measured by nanodrop (Thermo Fisher Scientific) and Agilent 2100 Bioanalyzer (Agilent).

### 5.9 Library Preparation

Preparation of the library was done by the Genomics and Bioinformatics Shared Resource (GBSR) at the University of Hawaii Cancer Center. In accoradnce with manufacturer’s instructions, QIAseq miRNA Library Kits (QIAGEN) and QIAseq miRNA NGS 12 Index IL (QIAGEN) were used for library construction. Library quality was assessed using the Agilent 2100 Bioanalyzer with a high-sensitivity DNA chip and sequenced using the Illumina NextSeq 500 at GBSR.

## Additional Files

### Additional File 1

#### Summary Report

Example report generated from the pipeline.

### Additional File 2

#### Rediscovery of 5’ and 3’ processing trends and 3’ D Box Motif in small RNAs

Here we delve deeper into the outputs of the pipeline in our example runs, showing that we are able to reproduce the findings of past authors.

### Additional File 3

#### Packaged normalization methods

CPM, TPM with fragment and miRNA counts, and ratio normalization methods to retain biological signal is discussed in an unsupervised clustering problem.

### Additional File 4

#### Additional applications

Additional applications regarding merging annotation databases are discussed in this supplementary file to highlight the need of choosing a reliable annotation database prior to utilizing the sRNAfrag pipeline.

### Additional File 5

#### Peak Calling Algorithm Implementation

Practical implementation of the peak calling algorithm.

### Additional Figure 1

#### Deviation from miRbase Loci

Called peaks are represented with red asterisk. Figures are generated through standard pipeline workflow. A) False peak for fragment derived from hsa-mir-500b with read coverage only present at loci that deviate from miRbase loci. B) Cluster region for predicted miRNA from hsa-mir-500b. C) 5’ and 3’ mature miRNA derived from hsa-mir-221 with peaks that deviate from miRbase loci. D) Cluster regions for miRNA predicted from hsa-mir-221.

### Additional Figure 2

#### Utilized Alignment Benchmark

Linear models depicting performance of AASRA method against synthetic sequencing data generated with the R Polyester package for multiple biotypes of sRNAs. Obtains a precision of 0.91 and a sensitivity of 0.95.

## Declarations

### Availability of data and materials

Cell line data generated from this study is available at GSE239311.

## Authors’ contributions

KN prepared the initial manuscript and developed the pipeline. MJ and MN shared preliminary data and thoughts that inspired development. MJ maintained/prepared cell lines. VK and MJ monitored the development of the pipeline from conceptualization to production. MH and YD assisted with conceptualization. All authors contributed to the final version of the manuscript.

## Acknowledgments

We would like to thank Dr. Karolina Peplowska of the GBSR at the UHCC for her excellent small RNA-seq performance. We would like to thank the Bank of Hawaii for facilitating the research funding from the Chun Foundation and the Jean Epstein Foundation. The technical support and advanced computing resources from University of Hawaii Information Technology Services – Cyberinfrastructure, funded in part by the National Science Foundation CC* awards # 2201428 and # 2232862 are gratefully acknowledged.

## Code availability

Our repository can be accessed at https://github.com/kenminsoo/sRNAfrag, DOI: 10.5281/ZENODO.8264987.

