## Additional File 1 for "sRNAfrag: A pipeline and suite of tools to analyze fragmentation in small RNA sequencing data"

### sRNAfrag Summary Report

#### P1 Module Outputs

No figures are produced during the first module. Primary function is to align and obtain counts.

##### Directory Structure

Main-Dir

|__out

```
|__tables     |__num_reads_bam.csv // Number of reads for each library. Can be used for CPM/TPM/RPKM
normalization.
    |__sRNA_frag_counts.csv // Counts for all fragments. Raw data.
```

#### S1 Module Outputs

Basic analyses and information regarding transcripts. Prepares for P2 module.

##### Count Distribution and Correction

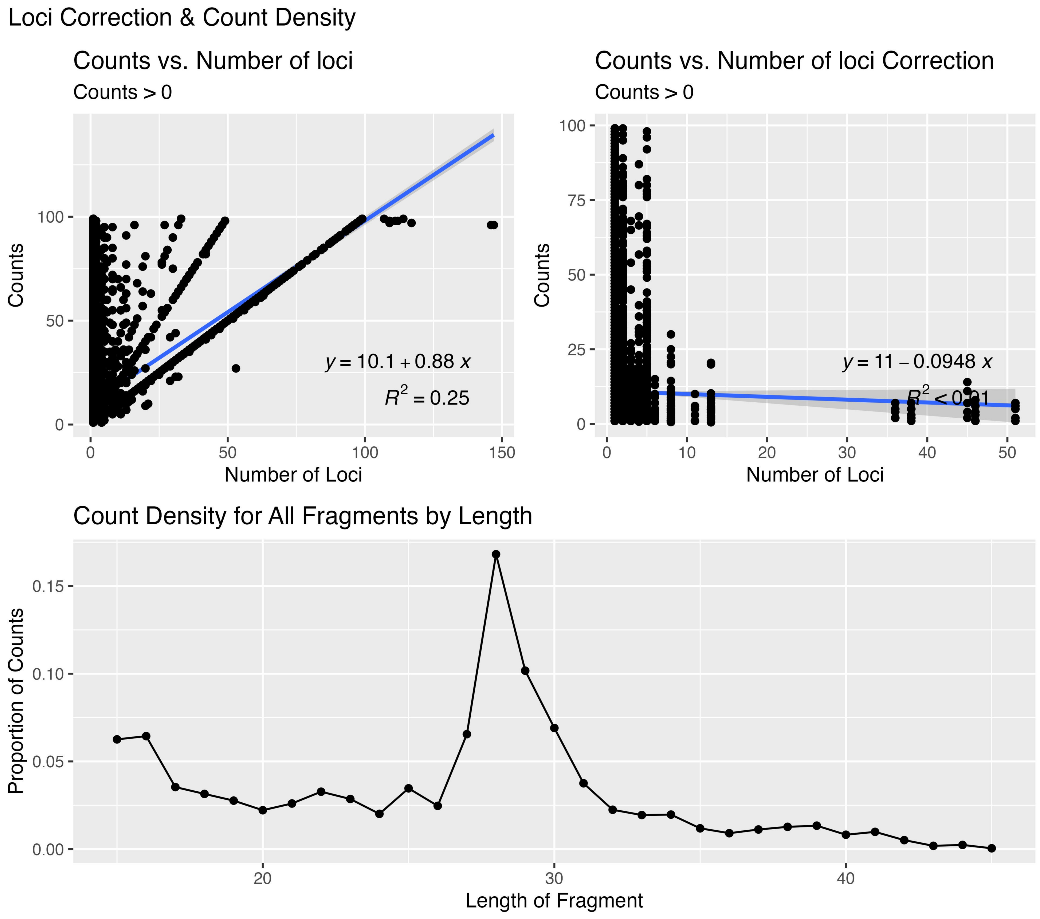


Figure 1. Pre and post correction counts with linear fits. Count density is also depicted.

### Standardized Loci Location

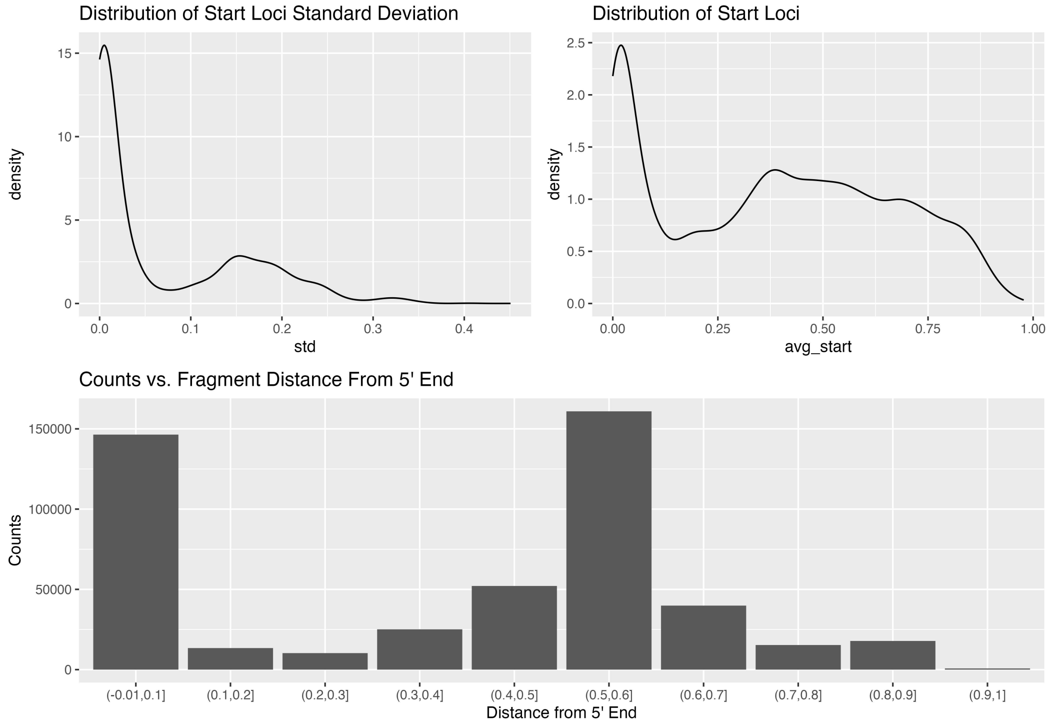


Figure 2. Start loci are standardized onto a 0-1 scale. If start loci do not agree (due to a fragment possibly being derived from multiple sources) its average is taken and the standard deviation is plotted. The average standardized loci is plotted by frequency. Finally, the count distribution is binned and plotted using a bar graph.

### Motifs by Frequency

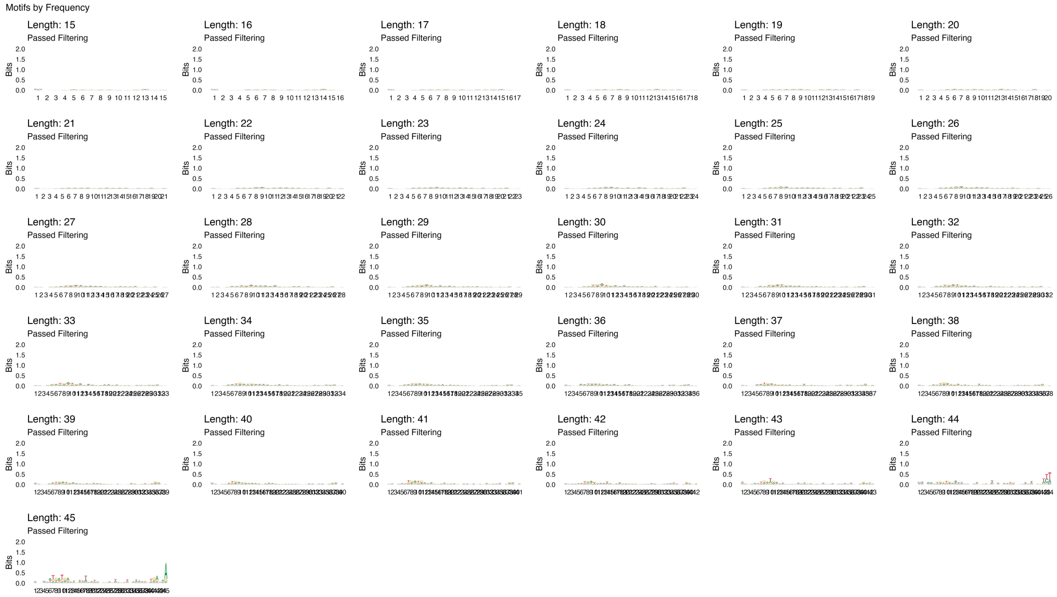


Figure 3. Logos of most common motifs by frequency.

#### Motifs by Counts

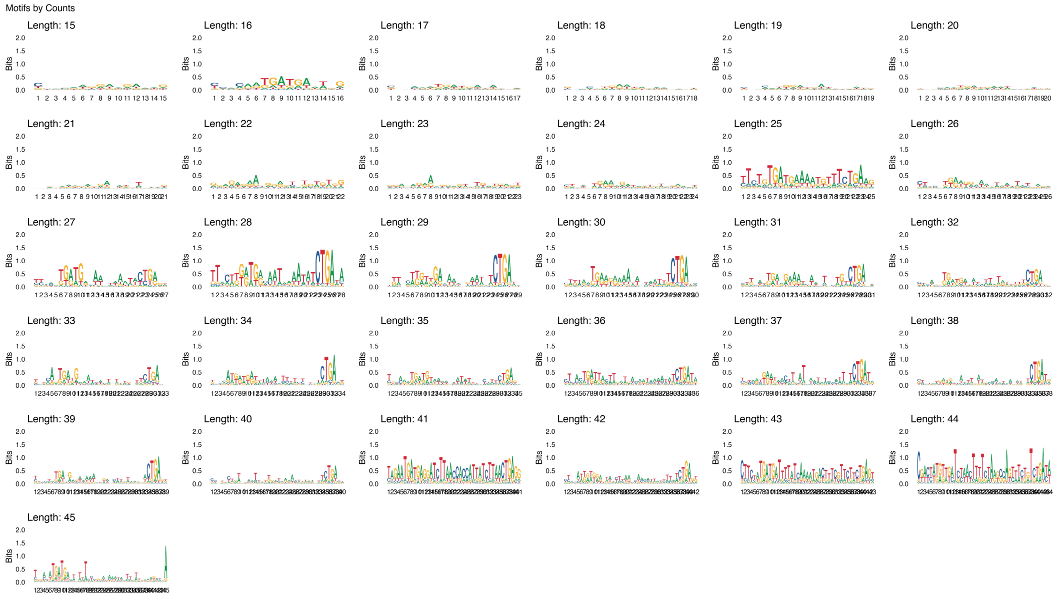


Figure 4. Logos of most common motifs by counts.

#### Most common 5p 5-mers

**Top 10 5' Flanking 5-mers**

| flanking5p | n | prop |
|---|---|---|
| CTGAA | 191 | 0.006248569 |
| TGAAA | 155 | 0.005070828 |
| TCTCT | 143 | 0.004678248 |
| TCTGA | 138 | 0.004514673 |
| GCTGG | 134 | 0.004383813 |
| AAACC | 133 | 0.004351098 |
| ATGAT | 133 | 0.004351098 |
| CTGAG | 133 | 0.004351098 |
| TTTCT | 127 | 0.004154807 |
| TGATG | 121 | 0.003958517 |

Figure 5. Most common fiver-mers on 5p end of fragment.

##### Most common 3p 5-mers

| Top 10 3' Flanking 5-mers | | |
|---|---|---|
| flanking3p | n | prop |
| AAAAA | 145 | 0.004743678 |
| CTGAG | 142 | 0.004645533 |
| TCTGA | 140 | 0.004580103 |
| TGCCA | 119 | 0.003893087 |
| TTCTG | 117 | 0.003827657 |
| AAAAT | 114 | 0.003729512 |
| TGAAA | 110 | 0.003598652 |
| CTGAA | 107 | 0.003500507 |
| CTGAT | 105 | 0.003435077 |
| TTTTC | 88 | 0.002878922 |

Figure 6. Most common fiver-mers on 3p end of fragment.

#### Directory Structure

Main-Dir

|__out

```
|__tables
|   |__num_reads_bam.csv // Number of reads for each library, Can be used for CPM/TPM/RPKM normalization
|   |__sRNA_frag_counts.csv // Counts for all fragments, Raw data
|   |__filtered_corrected_counts.csv // Adjusted counts
|   |__filtered_counts.csv // Unadjusted counts
|
|__figures
    |__S1_correction_density.jpeg // Figure depicting adjusted and count distribution
    |__S1_freq_motifs.jpeg // Motifs by frequency
|__S1_motifs_counts.jpeg // Motifs by counts
    |__S1_std_loci.jpeg // Loci location information
    |__S1_five-mer_3p.png // Most common 3p fiver-mers | Please feel free to modify the S1_figures.R     |__S1_five-
mer_5p.png // Most common 5p fiver-mers | If you'd like to generate different length.
```

#### P2 Module Outputs

##### Number of Sources for Each Fragment

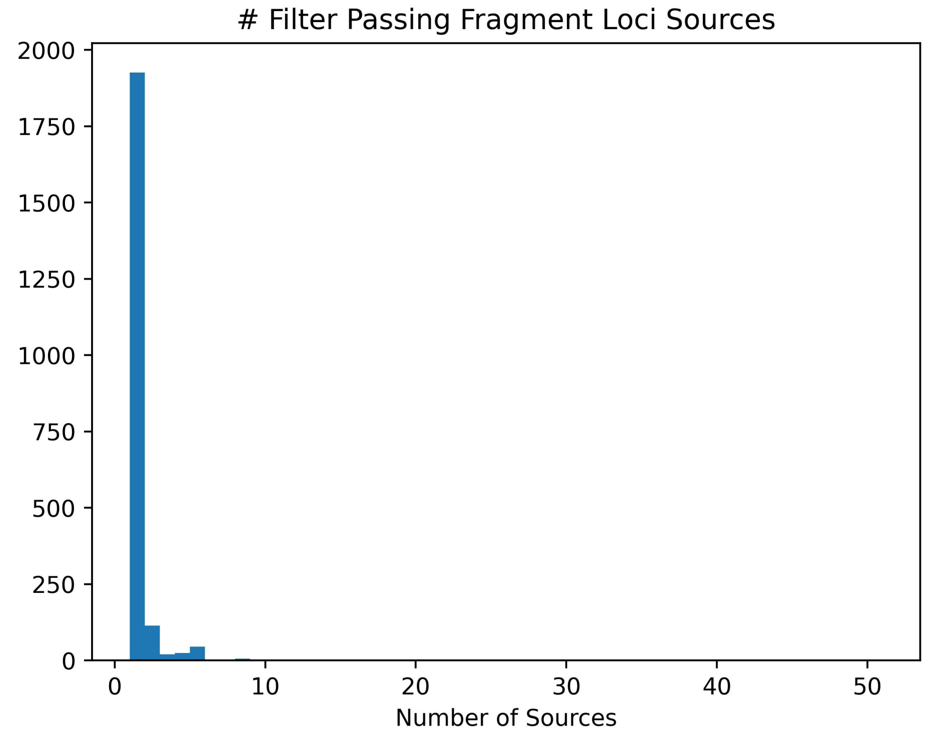


Figure 7. Distribution of number of potential sources each fragment has.

##### Number of Transcripts that span two clusters

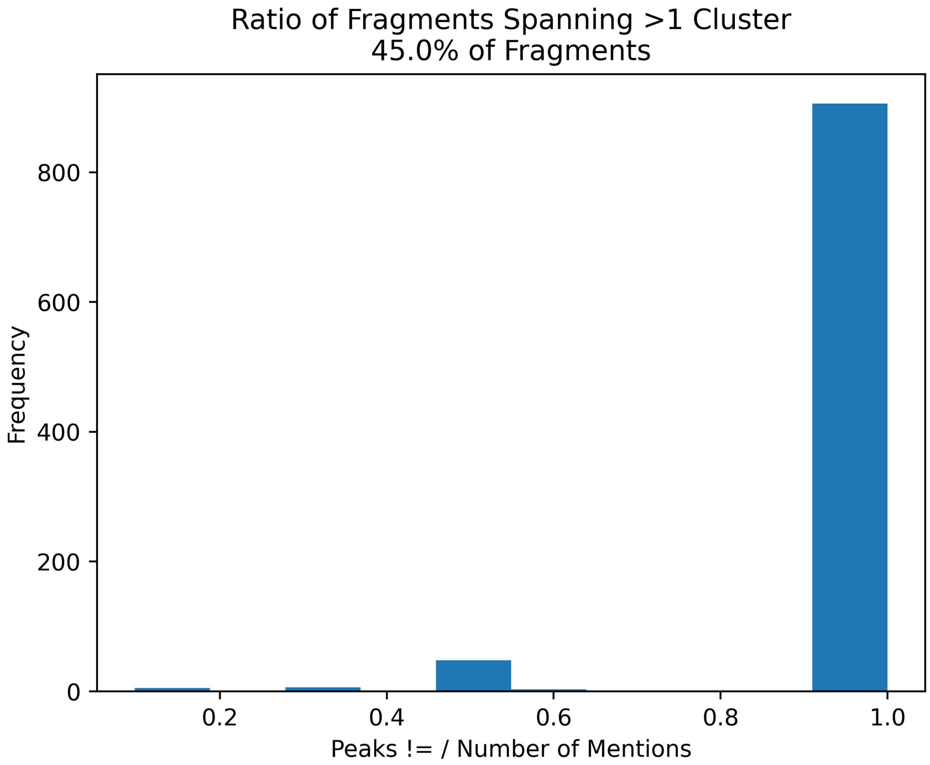


Figure 8. Ratio of disagreements (i.e. start = cluster 1, end = cluster 2) over total mentions. May signify interesting fragmentation pattern.

#### Directory Structure

Main-Dir

|__out

```
|__tables
|   |__num_reads_bam.csv // Number of reads for each library, Can be used for CPM/TPM/RPKM normalization
|   |__sRNA_frag_counts.csv // Counts for all fragments, Raw data
|   |__filtered_corrected_counts.csv // Adjusted counts |   |__filtered_counts.csv // Unadjusted counts |
|__cluster_peak_relationship_table.csv // Associates each start and end loci with a count and if it is a primary peak.
|   |__alternate_peaks.csv // Marks peaks that span two peaks. Used for plots in cluster folder.
|   |__annotated_ref_table.csv // Reference table associating fragment ID with merged ID. Includes information about external maps.
|   |__merged_counts.csv // The final output with sample counts and the set of license plates and external maps. |
|__figures
|   |__S1_correction_density.jpeg // Figure depicting adjusted and count distribution
|   |__S1_freq_motifs.jpeg // Motifs by frequency
|   |__S1_motifs_counts.jpeg // Motifs by counts
|   |__S1_std_loci.jpeg // Loci location information
|   |__S1_five-mer_3p.png // Most common 3p fiver-mers | Please feel free to modify the S1_figures.R
|   |__S1_five-mer_5p.png // Most common 5p fiver-mers | If you'd like to generate different length.
|   |__P2_Filter_Passing_Hist.jpeg // The number of sources that fragments could have. Distribution.
|   |__P2_cluster_start_end_disagreement.jpeg // Distribution of situations where start and end clusters are not equal. Not necessarily bad. P
|
|__sequences |   |__filtered_sequences.fa // Sequences in the fasta format. Do Gibbs Sampling to find motif. Or do
target prediction.
| |__source_peaks // If makelots is true, each source will have counts plotted against loci
position. |   |__source1
|   |__source2
|   |__... |
|__source_n
| |__cluster // If makelots is true, each source will have the cluster zones against loci highlighted, along with disagreements
plotted. |   |__source1
|   |__source2 |   |__... |   |__source_n | |__int_files // Intermediate script files pasted into the directory. Please take a look if
you'd like to contribute on the github repo!
    |__scripts...
    |__extra_files...
```
