## Additional File 2 for "sRNAfrag: A pipeline and suite of tools to analyze fragmentation in small RNA sequencing data"

**S2 File – Rediscovery of Fragmentation Patterns**

Main:

Li et. al. reported in 2012, “Extensive terminal and asymmetric processing of small RNAs from rRNAs, snoRNAs, snRNAs, and tRNAs.” Here we evaluate one statement in their paper.

“All ncRNAs generate both 5′ and 3′ products except snRNAs”

M. musculus snoRNA:

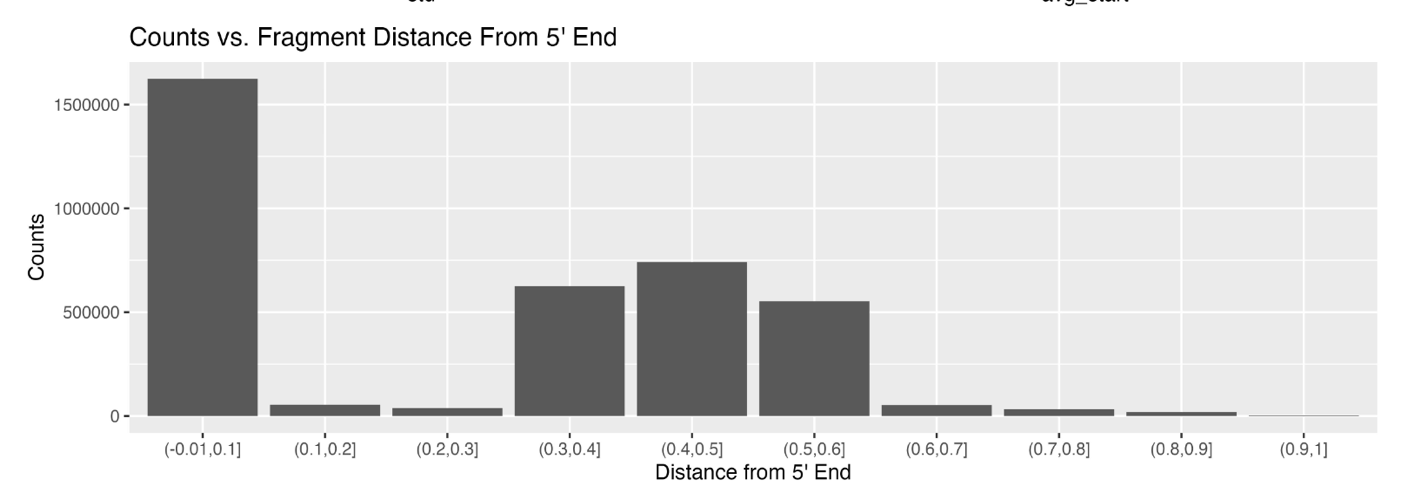

C. elegans snoRNA:

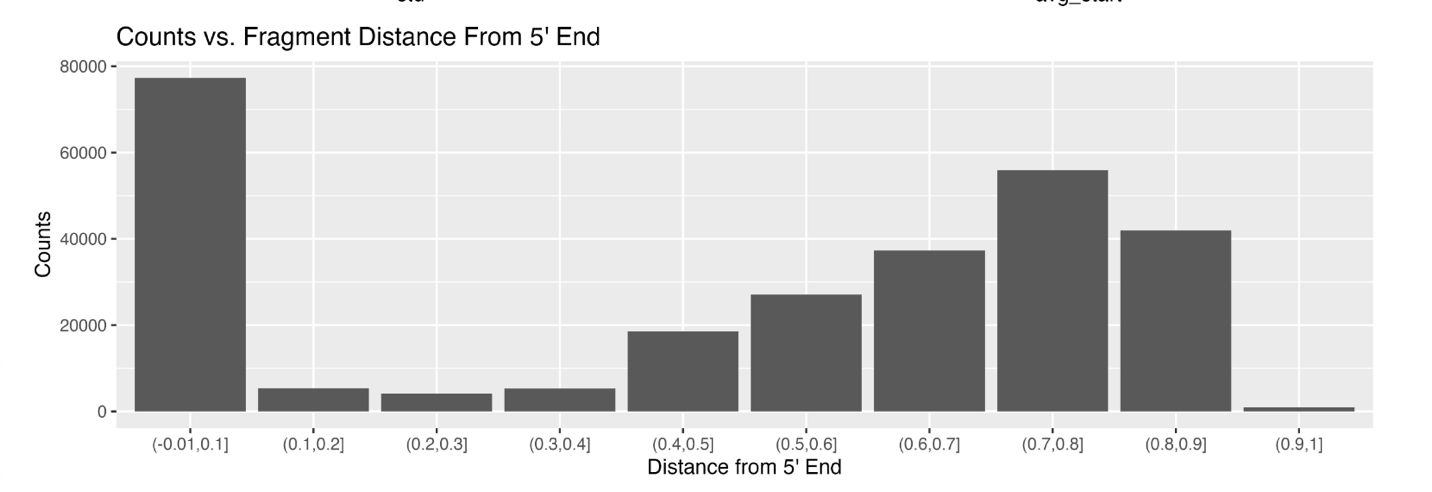
A. thaliana snoRNA:

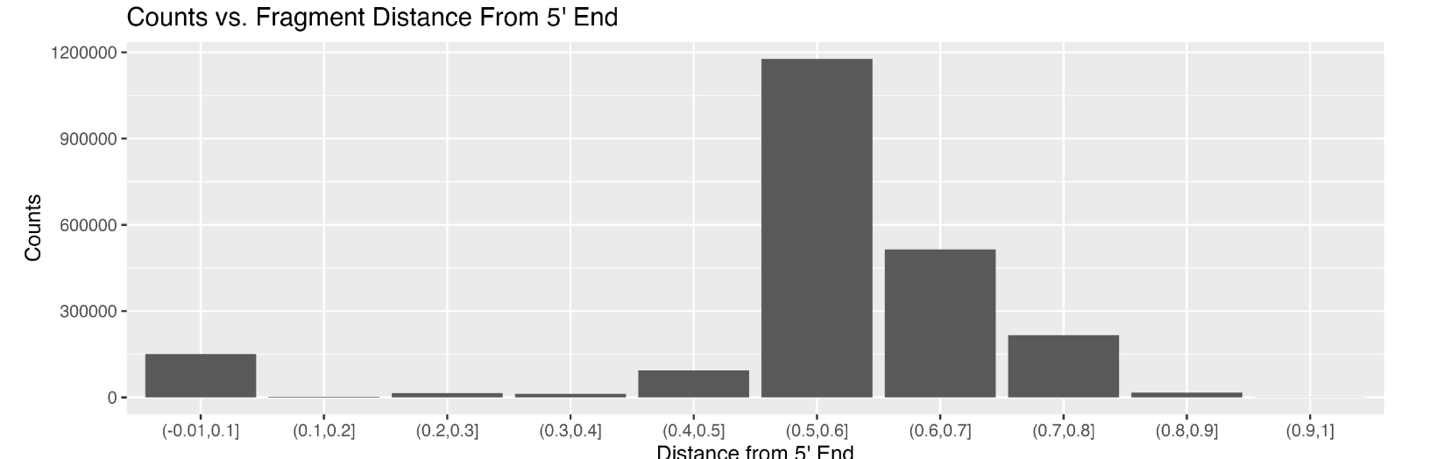

H. sapiens snoRNA:

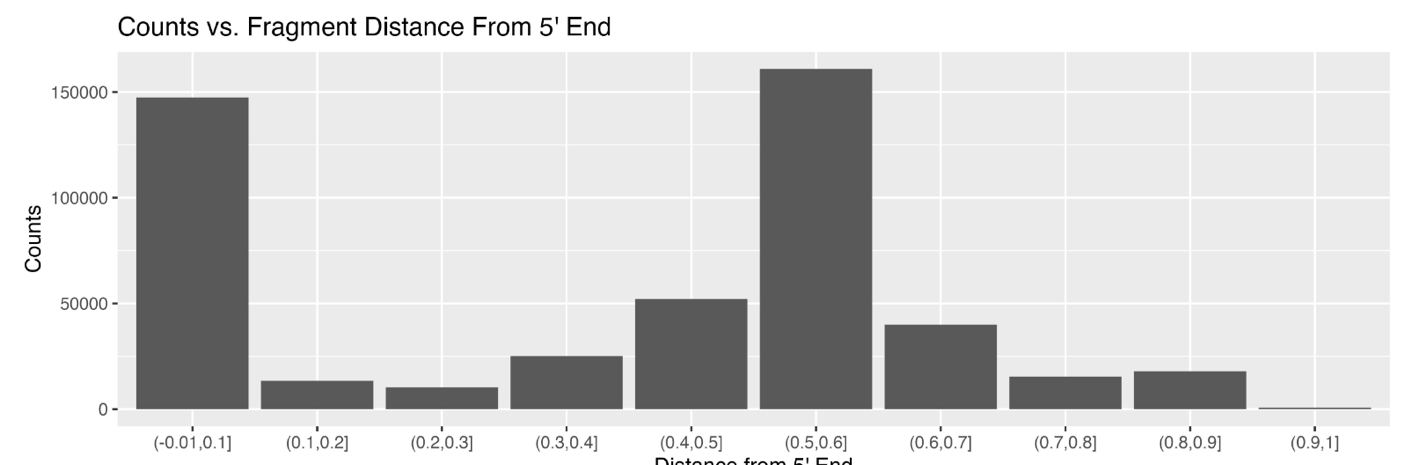

M. musculus rRNA:

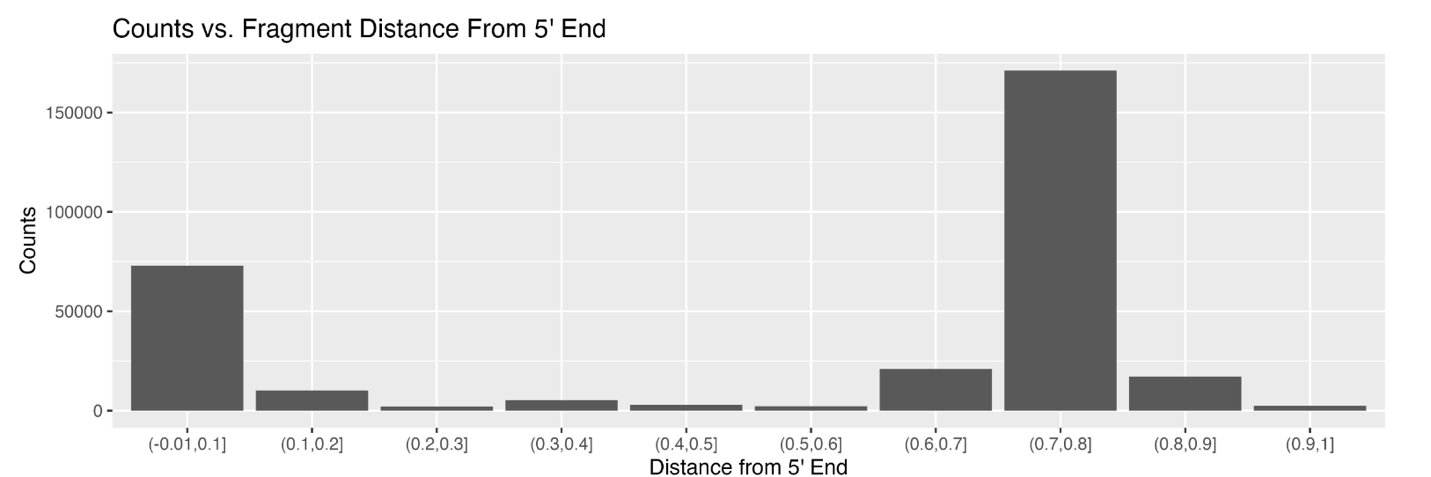

C. elegans rRNA:

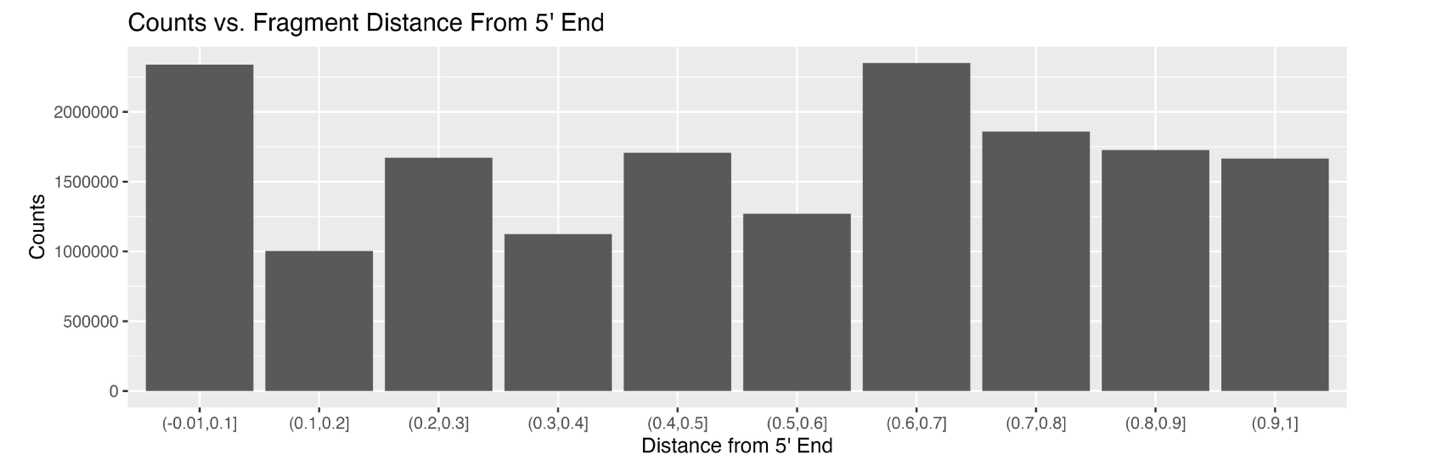

Technical variation?

A. thaliana rRNA:

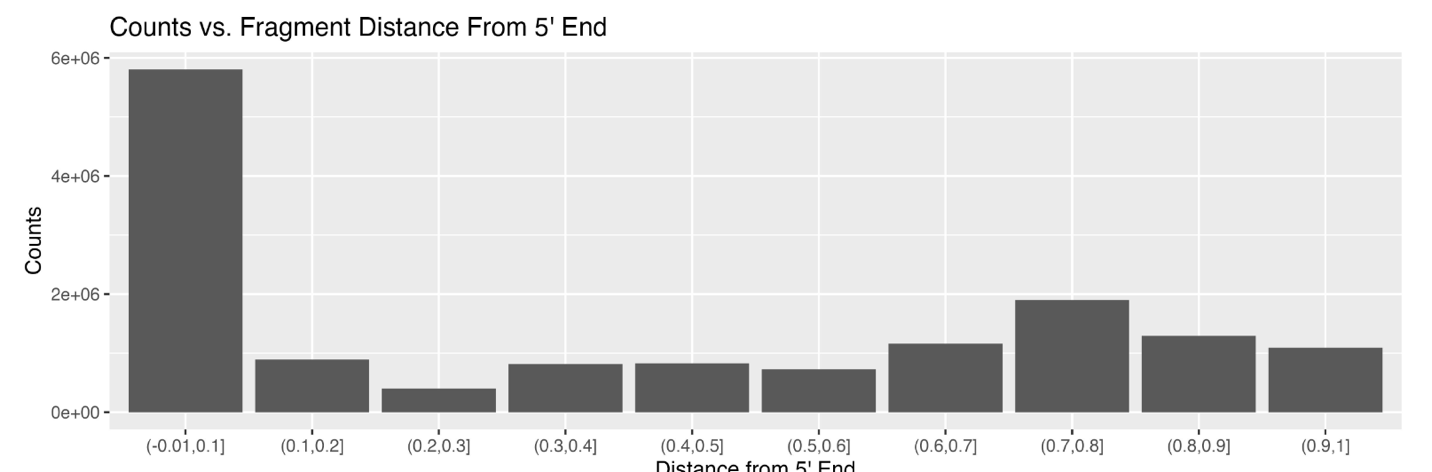

H. sapiens rRNA:

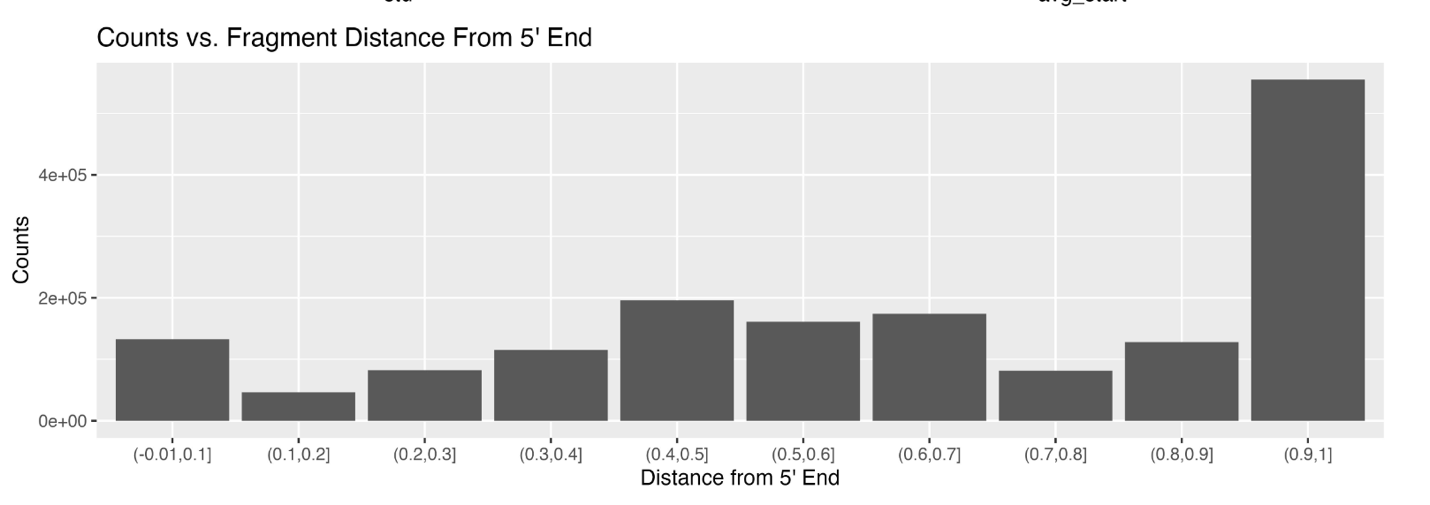

M. musculus snRNA:

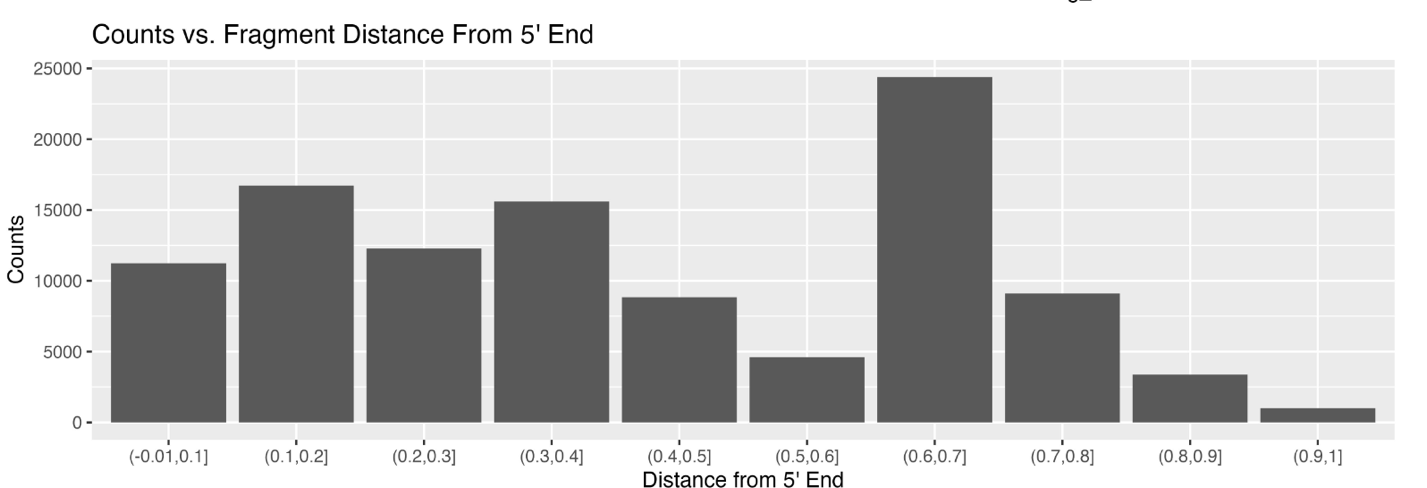

Preference for 3’, but not exclusive.

C. elegans snRNA:

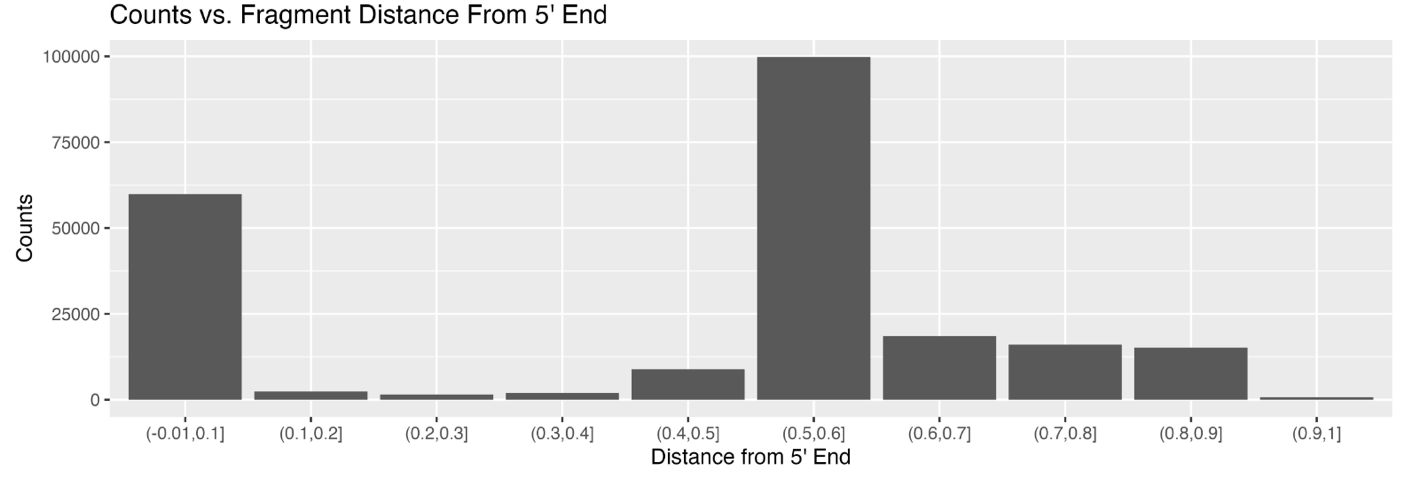

Preference for 3’, but not exclusive.

A. thaliana snRNA:

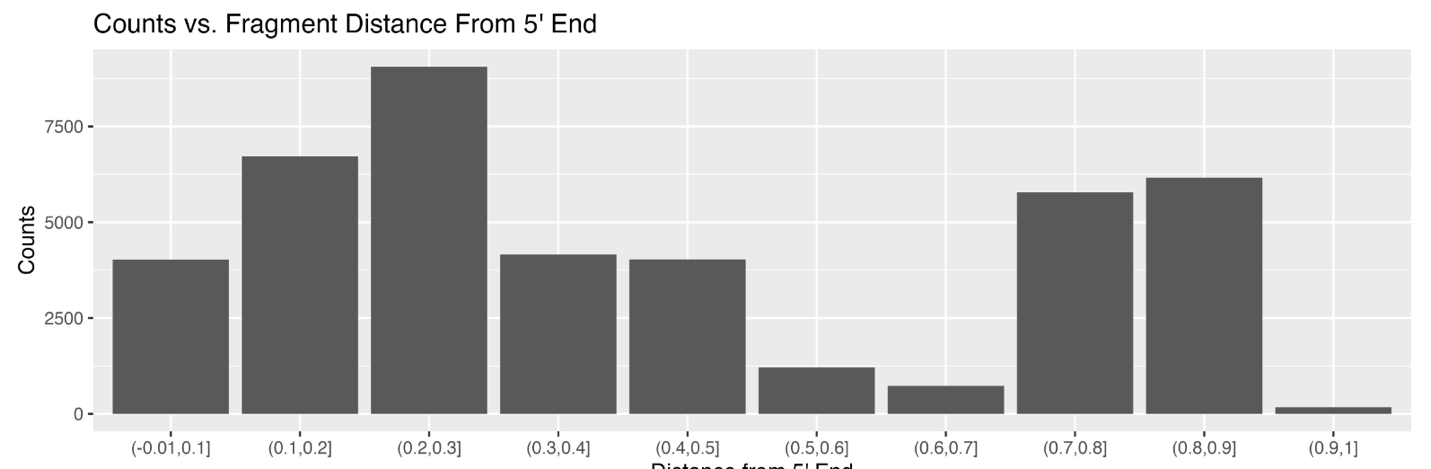

Plant is only one without 3’ preference.

H. sapiens snRNA:

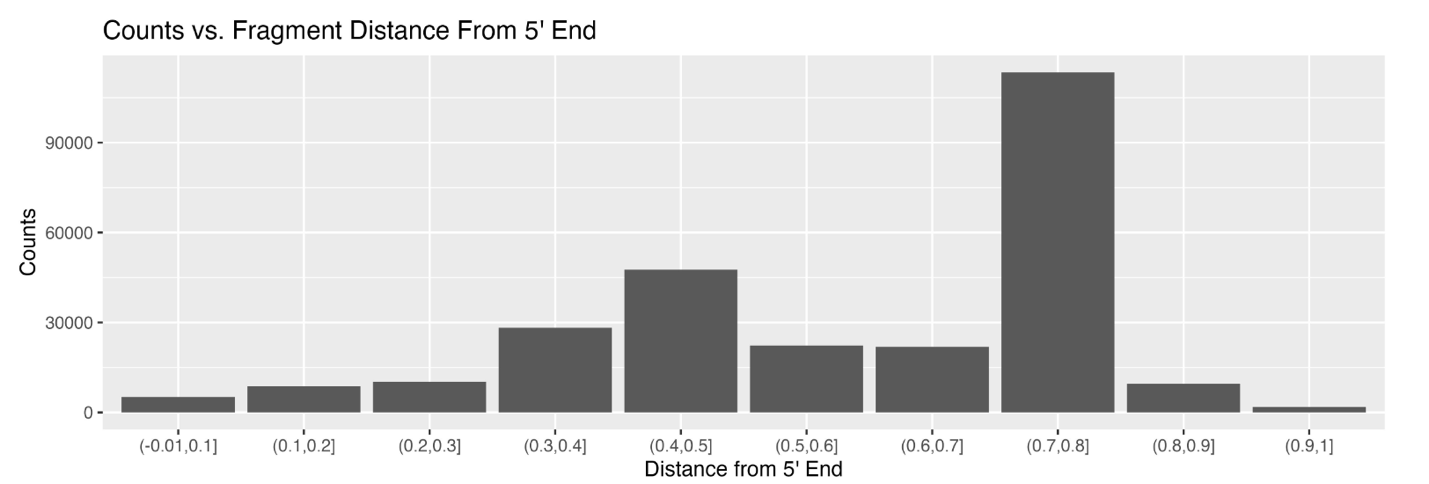

Here, we can see that small ncRNAs are terminally processed from the 5’ and 3’ ends of the transcript. Interestingly, A. thaliana was the only species to not exhibit 3’ preference in the processing of snRNAs.

Note, D box motif is CTGA.

Scott et. al. reported in 2012 that “Human box C/D snoRNA processing conservation across multiple cell types.” They specifically mention conservation of D box motif (54% of snoRNA derived fragments).

Mouse motifs, snoRNA, by counts:

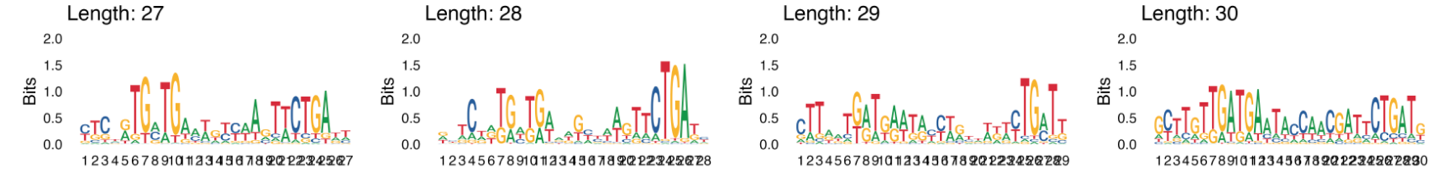

Aridopsis motifs, snoRNA, by counts:

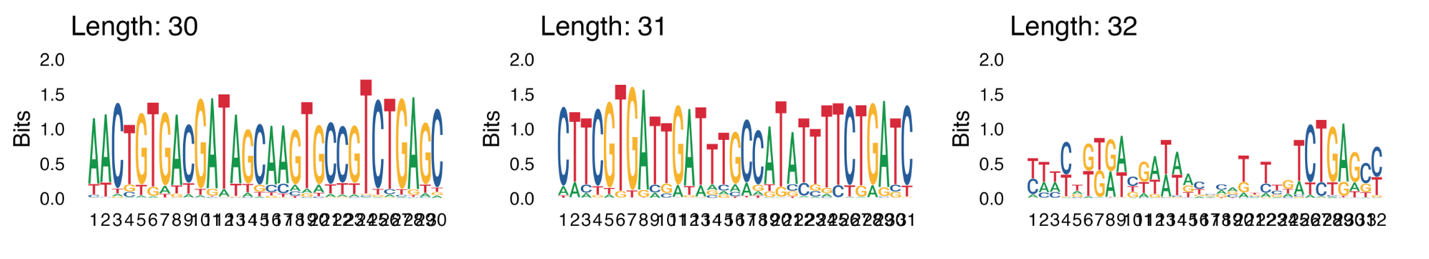

Human motifs, snoRNA, by counts:

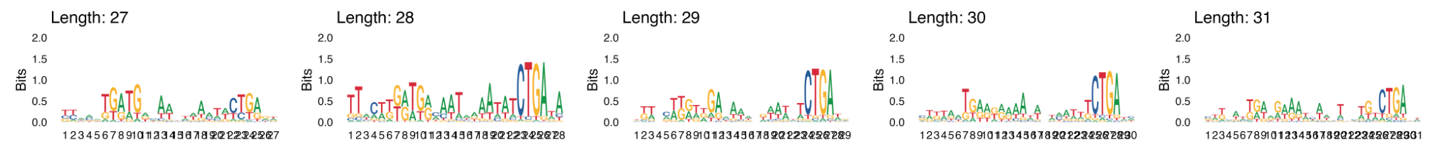

In each case, we see that the D box motif is conserved at the 3’ end of the fragment, typically 1-2 nucleotides away from the end of the fragment. Taft et. al. (2009) report that D box snoRNA derived fragments are primarily derived from the 3’ end of the snoRNA. This raises the question if there is a potential mechanism that allows the 3’ end of the snoRNA fragment to resist exonuclease activity.

This supplement works to show how the default outputs of sRNAfrag can be quickly looked at to identify patterns that may be of biological significance.
