## Additional File 3 for "sRNAfrag: A pipeline and suite of tools to analyze fragmentation in small RNA sequencing data"

**S3 File – Normalization Methods & Biological Significance**

Main Text:

One of the disadvantages of a pipeline that outputs non-standard tables such as ours, is that standard practices (for example, normalization) become more tedious as scripts must be adapted to accommodate. Thus, we have created built-in scripts to normalize using three methods: TPM with the fragment counts + mature miRNA counts (to ensure that it doesn’t appear overly high), the ratio of fragment counts to the sum of mature miRNAs with median >0, and finally the standard CPM, utilizing total reads. Here, we show that each does a valid job in maintaining the biological structure of the data by normalizing snoRNA fragments and attempt an unsupervised machine learning problem of clustering similar cell lines.

**TPM Normalization:**

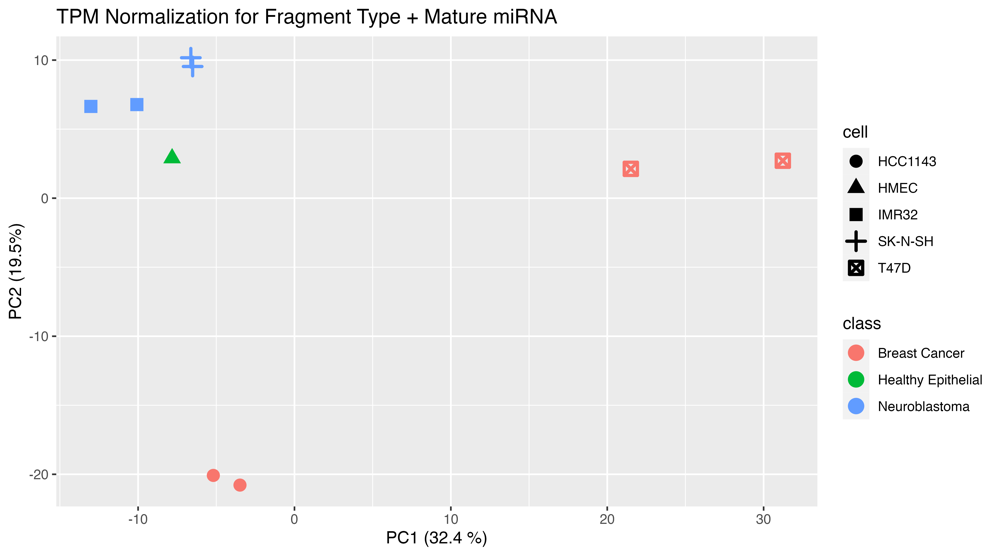

**Figure 1 PCA Plot for TPM Normalization.** Percentage indicates percentage of variance explained.

TPM normalization takes a slightly different approach by scaling the counts relative to the length of the RNAs and then normalizing to sum to one million. The logic here is that longer RNAs will, in theory, produce more fragments than shorter RNAs given equal expression levels. By considering both fragment counts and miRNA counts, TPM provides a balance between the abundance of small RNA fragments and their corresponding mature miRNAs. This method can be especially pertinent when the size distribution of RNAs varies across samples, ensuring that the observed differences are due to genuine changes in RNA expression and not merely due to differences in RNA length. However, it assumes a linear relationship between RNA length and fragment count, which might not always hold, especially in the context of fragmentation.

**Ratio Normalization**

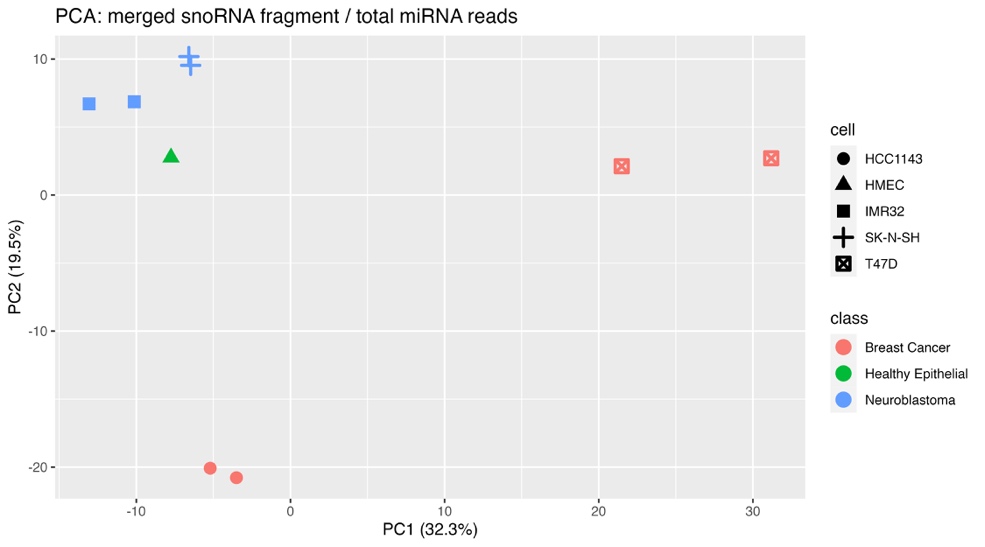

**Figure 2 PCA Plot for ratio normalization.** Percentage indicates percentage of variance explained.

This normalization technique emphasizes the relative abundance of the small RNA fragment compared to its mature miRNA counterpart. By tracking this ratio, insights into the processing and maturation pathway of the small RNA can be derived. But its effectiveness hinges on the accurate detection and quantification of the mature miRNA. If the mature miRNA is undetectable or its levels are erratic across samples, the normalization might yield skewed interpretations. Large variances in mature miRNA levels across samples could also obscure real biological differences in fragment expression. Note that the structure of the plots between ratio and TPM are very similar. Since they both essentially compare the relative amounts of fragment and miRNA, this is not exactly surprising.

**Counts Per Million Normalization**

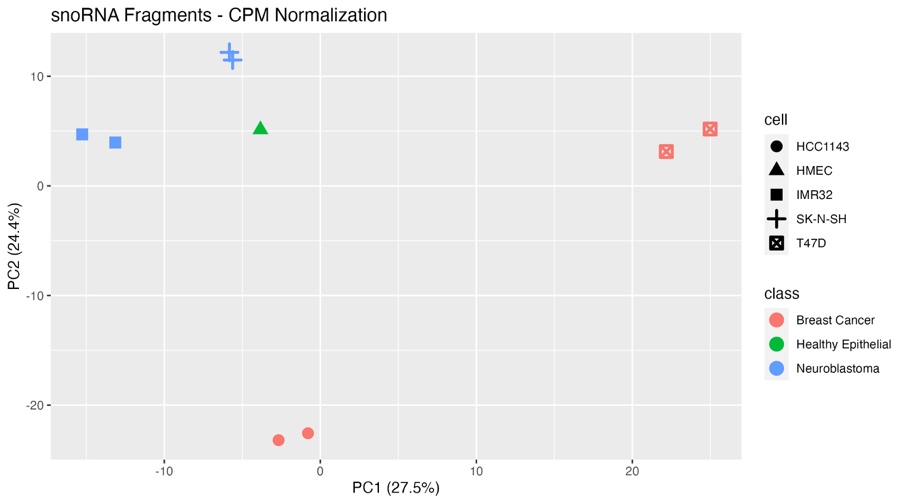

**Figure 3 PCA plot for CPM normalization.** Percentage indicates percentage of variance explained.

The CPM method in our study scales the raw count data by dividing each read count by the total number of sequencing reads (including both annotated and non-annotated reads) and then multiplying by one million. This normalization process ensures that the data is adjusted for the total sequencing output rather than just the subset of annotated reads. By converting raw counts into CPM in this manner, we account for any potential variations introduced due to differences in the overall sequencing depth across samples. This allows for more transparent comparisons of small RNA fragment abundance across various datasets. While CPM provides a snapshot of RNA abundance relative to the entirety of the sequencing output, it should be noted that this method does not correct for library composition effects. Thus, in samples where certain small RNAs are highly expressed, they might overshadow those with lower expression. In essence, while CPM offers a clearer perspective on RNA abundance given the entire sequencing landscape, it may still face challenges in capturing genuine biological variations when certain RNA populations are highly variable or when there are outliers.

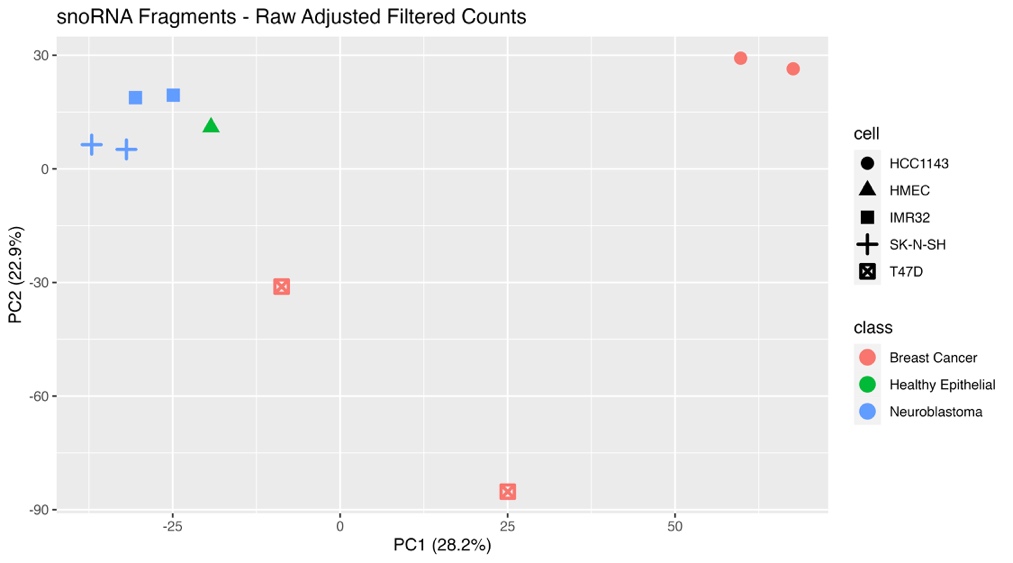

**Figure 4 PCA plot for filtered raw counts.** Percentage indicates percentage of variance explained.

Using raw counts provides a pure method of gauging small RNA fragment abundance. It offers an undistorted view of the RNA fragments' presence in the sample without any adjustments. This method is particularly valuable when samples are uniform in terms of sequencing depth and quality, giving a clear snapshot of the RNA environment. However, raw counts can be influenced by variations in sequencing depth. Any differences in library size or sequencing standards can introduce biases, blurring the line between genuine biological variations and technical inconsistencies. An important observation in our study was the consistent library size across all samples. Such uniformity means that many of the technical biases typically associated with raw counts are mitigated. Thus, we do not recommend using raw filtered counts.
