## Additional File 4 for "sRNAfrag: A pipeline and suite of tools to analyze fragmentation in small RNA sequencing data"

**S4 File.** Additional Applications

Main Text

The impact of annotation choice on downstream analyses is quite large. Even in bulk RNA-seq where annotations are quite well established, choosing which well established database to use can lead to differential analysis outcomes^1^. Authors have attempted to address this issue by generating new annotations^2^. However, such efforts are often futile as annotations continue to be updated. As a result, we were motivated to create scripts which can be accessed from the command line to generate and evaluate annotations. This supplementary file will show the intuition behind some of the tools packaged with the pipeline. It will show how commands packaged with the pipeline can be used to generate ensemble annotations from multiple sources. This is important as some fragments may be missed if full annotations are not utilized. Beyond this document, we recommend users consult the documentation on the github repository for readability and more information.

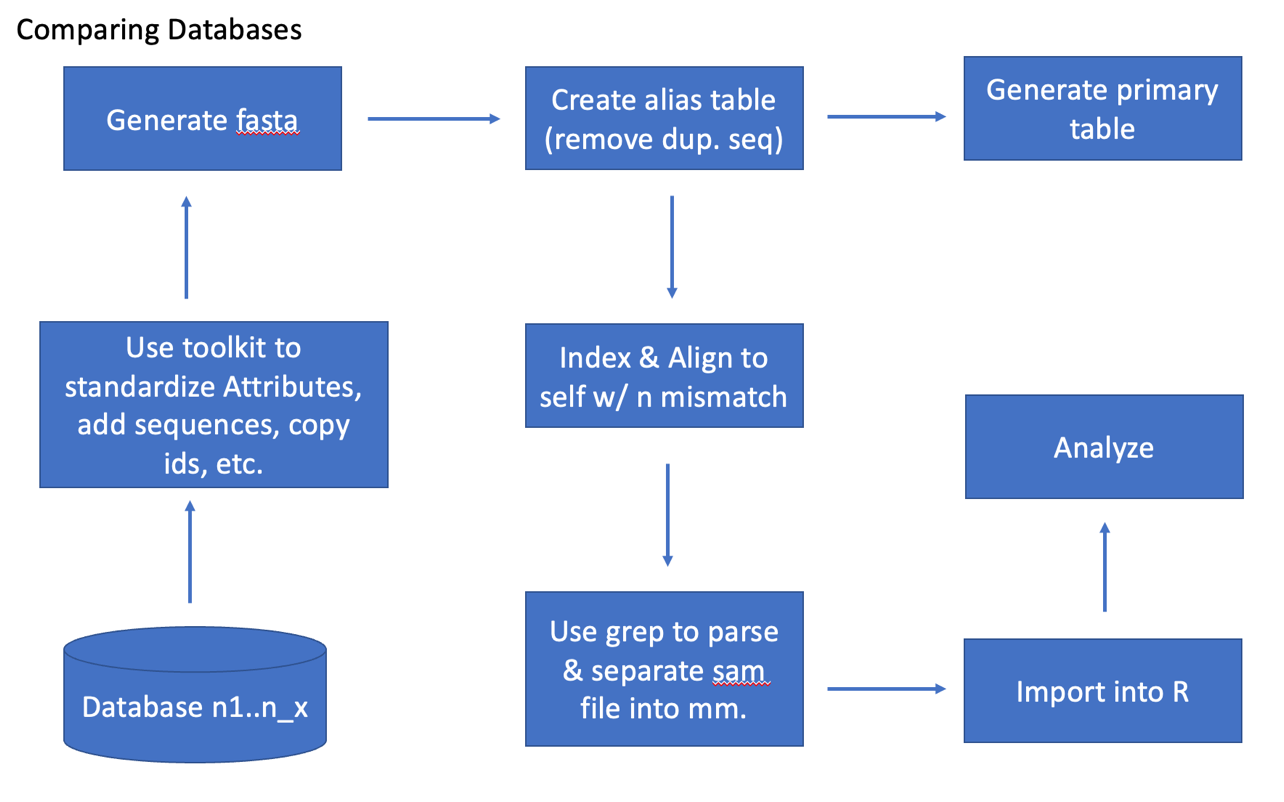

**S1 File, Fig 1. General workflow for combining databases.** Starting with multiple databases, tools packaged with the pipeline can be first used to standardize attributes between these different databases. Then, using unix commands, they should be joined into one file. Then a fasta of sequences is generated, with duplicates being marked and removed. An alignment with itself allows one to see how transcripts might map to each other and can be used when considering multi-mapping transcripts.

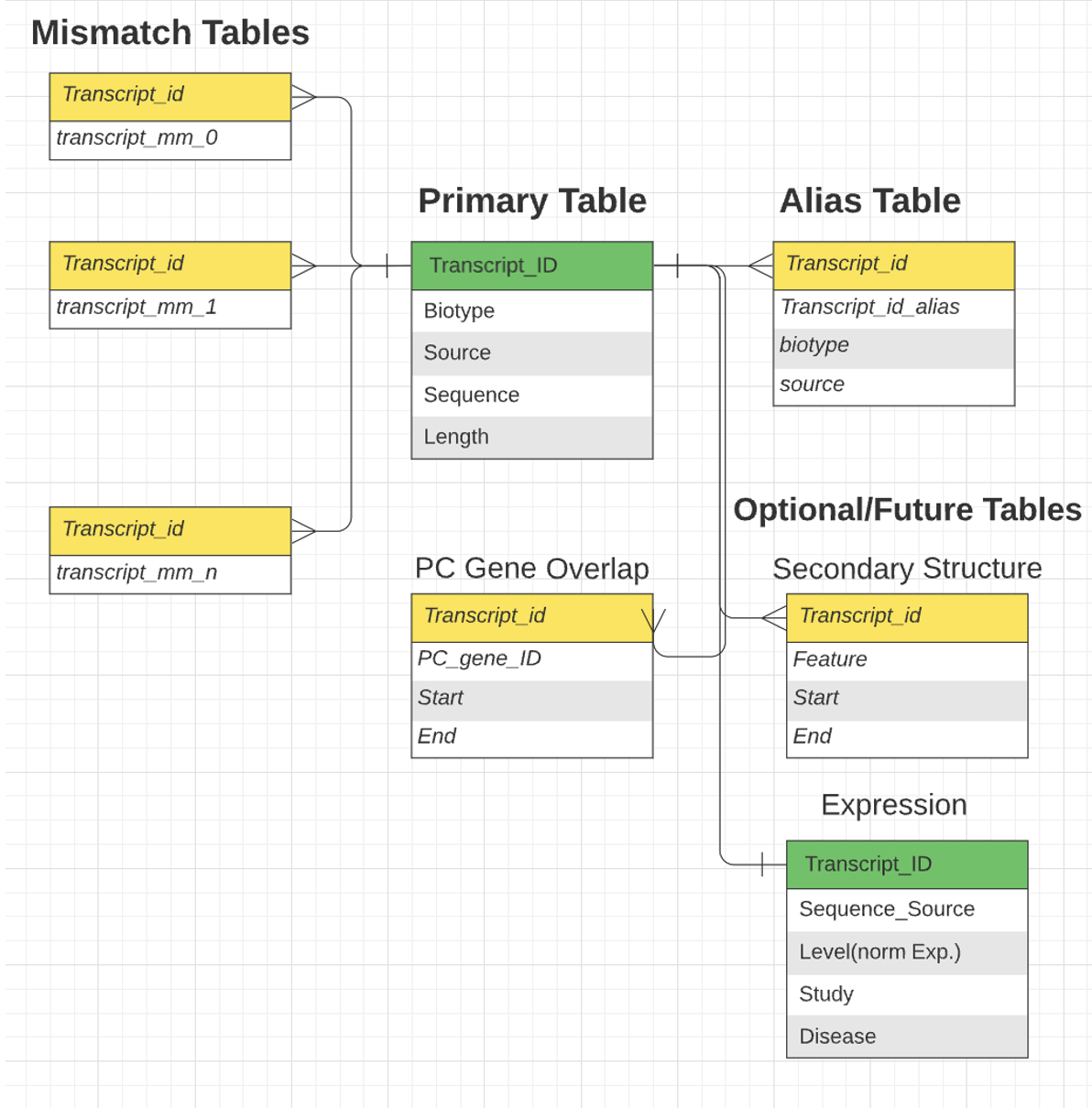

**S1 File, Fig 2. Structure of the Mismatch Tables.** A primary table with biotype, source, length, and sequence information is created with the merged databases. By maintaining transcript_ids (or any primary key), it allows users to connect it with other tables.

Protocol

1) Download databases of interest.
2) Standardize attributes and filter GTF if desired.
   a. Select based on column value
      i. python PATH/TO/gtf_modifiers.py select_column <**input_gtf>** <**out_name>** <**col_num** > <**value_to_select >**
   b. Select based on attribute value
      i. python PATH/TO/gtf_modifiers.py select_gtf <**input_gtf**> <**out_name**> <**attribute_name**> <**value_to_select**>
   c. Standardize attributes & select
      i. python PATH/TO/gtf_modifiers.py standardize_attributes <**input_gtf>** <**output_gtf**> <**mapping_dictionary**>
         1. {value_in_input:value_in_output}
         2. ‘{"transcript_id":"transcript_id", "transcript_biotype":"biotype"}'
            a. *Wrap {} in ‘’ & entries in “”
3) Combine GTF files
   a. cat gtf1.gtf gtf2.gtf … > gtf_new.gtf
4) Generate a fasta file
   a. python PATH/TO/conversion_tools.py gtf_to_fasta <**gtf_input**> <**output**> <**ref_genome**> <**primary_key**>
5) Create mismatch table and alias files.
   a. python PATH/TO/gtf_groundtruth.py generate_groundtruth <**gtf_input**> <**ref_genome**> <**output_dir**> <**primary_key**> <**information_dict**> <hisat = False> <num_mismatch = 2> <input_fasta = "sequences.fasta ">
      i. information_dict (dict): Dictionary specifying additional information to include in the primary table. The keys are the column names, and the values are lists in the format [gen_or_attribute, column_index_or_name], where gen_or_attribute is either 0 (for general data) or 1 (for attribute data), and column_index_or_name is the index or name of the column. i.e. {"biotype":[1, "biotype"], "source":[1, "databases"]}
      ii. The generated fasta file should be named sequences.fasta unless the user chooses to change it in this command.
6) Outputs
   a. Primary.csv // The primary table with data specified in information dict.
   b. Too_short.csv // Transcripts that were too short (<10 nt)
   c. Too_long.csv // Transcripts that were too long (>1000 nt)
   d. N_exist.csv // Transcripts that have N in their sequence
   e. Alias.csv // Transcripts with same name in the gtf file.
   f. n_mm.sam // All transcripts that aligned to other transcripts with n mismatches.
7) **OPTIONAL:** parse mismatch file to generated figures regarding mappings to other sequences.
8) Generate a new fasta file with only unique sequences
   a. python PATH/TO/conversion_tools.py tsv_to_fasta <**tsv**> <**output**> <**key_col**> <**seq_col**> <--delim=“,”>
9) Generate a new GTF file with the deduplicated fasta file.
   a. python PATH/TO/gtf_generation.py generate_from_fasta <**fasta**> <**output_gtf**> <**index_name**> <**new_dir**>
      i. Ensure to index a reference genome while generating.

We applied this ground truth generation to the RNA Central small RNA database for humans, allowing duplicate sequences to be collapsed **(S1 Table 1, S2 Table 2)**.

**S1 Table 1. Portion of the alias table**

| primary_transcript_id | alias_transcript_id |
| --- | --- |
| URS00004B825B_9606.190:ncRNA_exon1 | URS00004B825B_9606.1764:ncRNA_exon1 |
| URS00005E0C40_9606.191:ncRNA_exon1 | URS00005E0C40_9606.1765:ncRNA_exon1 |
| URS00000C5103_9606.193:ncRNA_exon1 | URS00000C5103_9606.1770:ncRNA_exon1 |
| URS000025781F_9606.196:ncRNA_exon1 | URS000025781F_9606.1772:ncRNA_exon1 |
| URS00006FBD25_9606.1773:ncRNA_exon1 | URS00006FBD25_9606.1773:ncRNA_exon1 |
| URS00025A9E25_9606.1781:ncRNA_exon1 | URS00025A9E25_9606.1781:ncRNA_exon1 |
| URS00025A9E25_9606.1781:ncRNA_exon2 | URS00025A9E25_9606.1781:ncRNA_exon2 |
| URS00025A9E25_9606.1781:ncRNA_exon3 | URS00025A9E25_9606.1781:ncRNA_exon3 |
| URS00025A9E25_9606.1781:ncRNA_exon4 | URS00025A9E25_9606.1781:ncRNA_exon4 |
| URS00005B3517_9606.202:ncRNA_exon1 | URS00005B3517_9606.1782:ncRNA_exon1 |
| URS00003B7688_9606.206:ncRNA_exon1 | URS00003B7688_9606.1790:ncRNA_exon1 |
| URS00002FCFC9_9606.213:ncRNA_exon1 | URS00002FCFC9_9606.1791:ncRNA_exon1 |
| URS0000034516_9606.1804:ncRNA_exon1 | URS0000034516_9606.1804:ncRNA_exon1 |

The alias table identifies both transcripts that appear multiple times in an annotation file and those that have the same sequence with different IDs. One example of each of these cases are marked in red and green, respectively.

**S1 Table 2. Summary of Generated Alias Table**

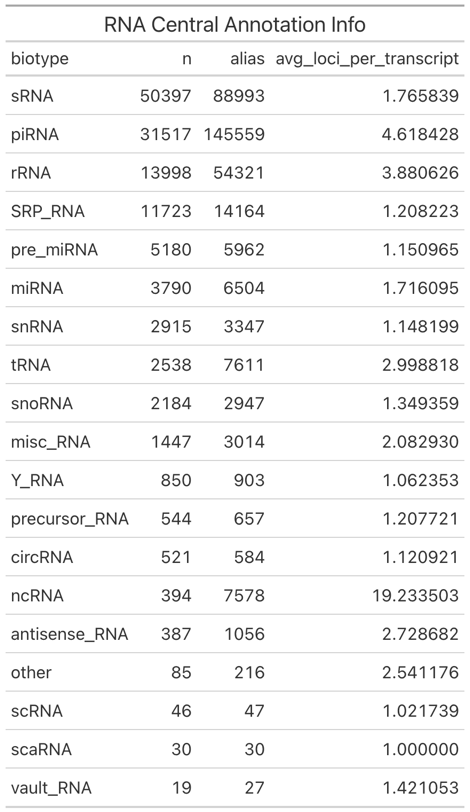

The alias table generated from RNA Central contains sRNAs (small RNAs) which often have aliases that map to other biotypes of small RNAs. RNA central can be thought of as one large, joined database. Our script parses these annotations, only keeping those that are unique, preserving their relationships in an external table.

Further analysis of the “n_mm.sam” files (which can be imported into R as tsv files after skipping the header), reveals that many small RNA sequences can be found within other small RNAs (S1 File, Fig 3.)

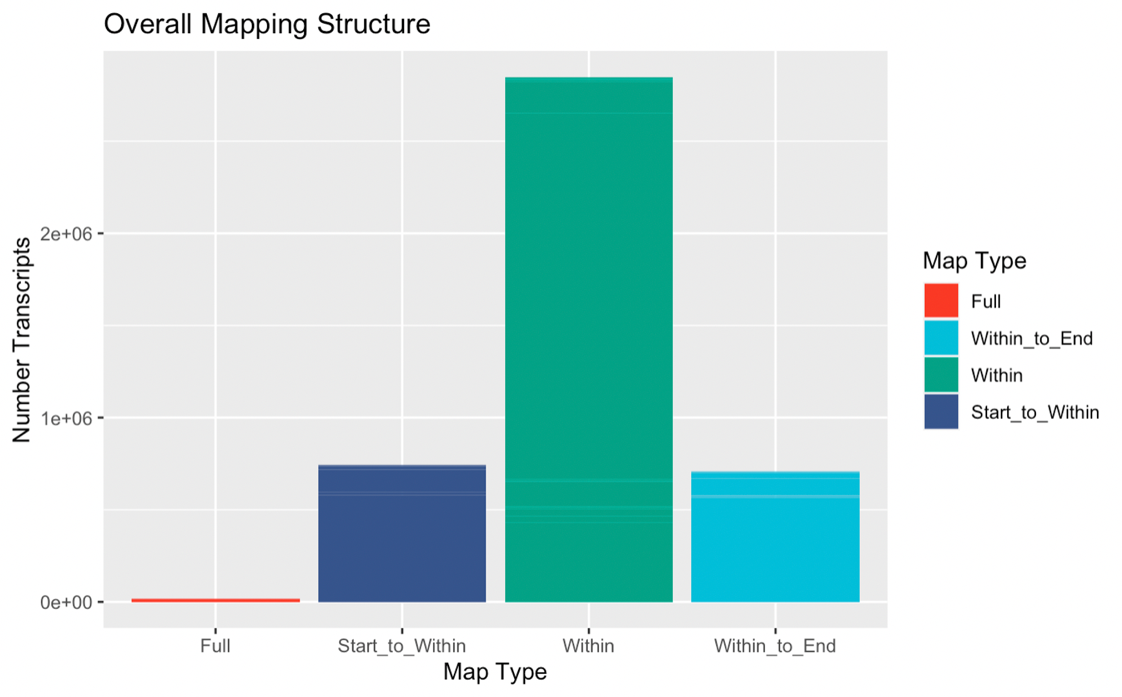

**S1 File Fig 3. Majority of transcripts map within other transcripts**

Within mappings refer to those that have start and end positions that do not correspond to the start or end positions of the transcript that was mapped to. This occurs for over 2 million transcripts within the RNA Central database.

Given the severity of the multi-mapping problem in the small RNA space, it is important to select carefully curated databases to ensure accurate downstream analyses. We also introduce one potential way to merge annotations from different databases, collapsing duplicated sequences while also preserving their original identities in an external table. However, this is not a new issue in biological databases. Chen et. al. articulates this problem extremely well^3^. We acknowledge that our scripts do not solve the data quality issue. We recognize that it represents a much larger, systemic issue with how biological databases are curated and published.
