## Additional File 5 for "sRNAfrag: A pipeline and suite of tools to analyze fragmentation in small RNA sequencing data"

**S5 File. Algorithm Implementation**

Main Text

It is certainly true that our clustering and peak calling algorithm could be implemented using standard loops, dictionaries, and lists. However, we decided to use Pandas and NetworkX to implement our algorithm. This supplemental will cover peak detection which was only covered in brief in our article.

Firstly, a graph is initialized with all sources connected to all fragments that it could have potentially derived from (S2 Fig 1.)

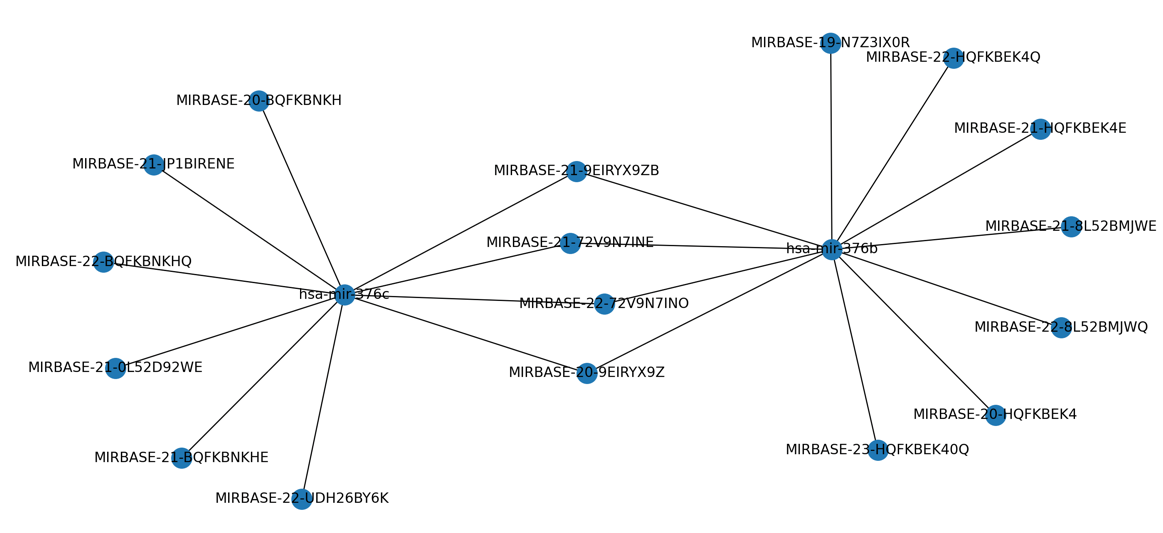

**S2 Fig 1 Initialized graphs**. The initial graph holds information regarding fragments that can be derived from two sources. This increases the complexity of the algorithm which led us to generate star graphs as opposed to simple bipartite graphs.

These graphs are bipartite graphs because sources only connect to fragments, and vice versa. This simplifies the algorithm a lot, allowing us to save computational time. From this graph, a star graph is created with the source as the center node (S2 Fig 2.)

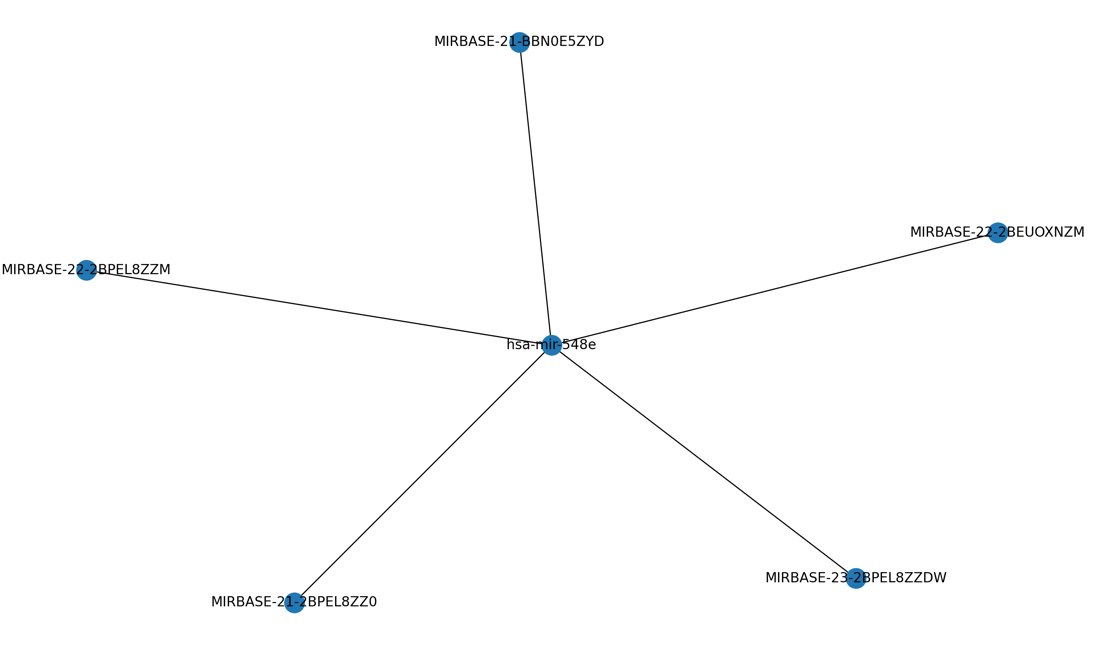

**S2 Fig 2 Star graphs.** Star graphs are generated with source transcript as the center node and fragments (that are filter passing) with edges to the center node.

The edges in this graph hold data regarding loci and counts. Counts are aggregated position wise, meaning all counts with the same positions are summed. A data frame is generated for each source beginning one nucleotide before from the smallest loci position (i.e. if 1, then 0; if 10, then 9). This data frame is used to detect peaks with the formulas seen in the paper (S2 Table 1.). Start loci and end loci are kept in memory as separate data frames. The example will be shown for only one type (i.e. start or end). NZ is the Boolean expression “counts > 0”. “_A” or “_B” represents a shifted column, either after (a shift of -1) or before (a shift of 1). ZSM refers to a zero count that is sandwiched between two non-zero values and is zero itself, represented as the expression “NZ_B == True & NZ_A == True & NZ == False.” This is to generate the “Add” column, a value that is added to counts to eliminate sandwiched zeroes. The add column is generated by multiplying the next counts by ZSM, a Boolean value that, when true, is treated as 1.

**S2 Table 1. Peak Detection Data Frame — Sandwiched 0 Elimination**

| Loci | Counts | NZ | NZ_B | NZ_A | ZSM | Counts_A | Add |
| --- | --- | --- | --- | --- | --- | --- | --- |
| 0 | 0 | F | NA | T | F | 20 | 0 |
| 1 | 20 | T | F | T | F | 40 | 0 |
| 2 | 40 | T | T | T | F | 70 | 0 |
| 3 | 70 | T | T | T | F | 50 | 0 |
| 4 | 60 | T | T | F | F | 0 | 0 |
| 5 | 0 => 10 | F | T | T | T | 10 | 10 |
| 6 | 10 | T | F | F | F | 0 | 0 |
| 7 | 0 | F | T | F | F | 0 | 0 |
| 8 | 0 | F | F | T | F | 30 | 30 |
| 9 | 30 | T | F | F | F | 0 | 0 |
| 10 | 0 | F | T | NA | F | NA | NA |

*The adjustment is in red.

Adjusted counts with eliminated sandwiches are then used to detect peaks. The new column with adjusted counts will be called “AdjC.” The only change that occurred is found in loci 5, where the sandwiched zero is eliminated. The next step is to calculate changes and to smooth out these changes to reduce the number of potential false positive peak calls (S2 Table 2.). First, the adjusted counts column is shifted one down (AdjC_B). Then, the ΔSC is calculated by subtracted AdjC_B from AdjC (AdjC – AdjC_B). The ΔSC is then shifted one row up (ΔSC_A). ΔSC_A * 0.5 + ΔSC is then calculated to be ΔSC_Mod. Two Boolean expressions are evaluated to generate ΔSC_Mod_+ (ΔSC_Mod > 0) and ΔSC_A_+ (ΔSC_A > 0) . A final Boolean expression is evaluated to determine if a loci is a peak (ΔSC_Mod_+ == True & ΔSC_A_+ == False).

**S2 Table 2. Peak Calling**

| Loci | AdjC | AdjC_B | ΔSC | ΔSC_A | ΔSC_Mod | ΔSC_Mod_+ | ΔSC_A_+ | Peak |
| --- | --- | --- | --- | --- | --- | --- | --- | --- |
| 0 | 0 | NA | NA | 20 | 10 | T | T | F |
| 1 | 20 | 0 | 20 | 20 | 30 | T | T | F |
| 2 | 40 | 20 | 20 | 30 | 35 | T | T | F |
| 3 | 70 | 40 | 30 | -10 | 25 | T | F | T |
| 4 | 60 | 70 | -10 | -50 | -35 | F | F | F |
| 5 | 10 | 60 | -50 | 0 | -50 | F | F | F |
| 6 | 10 | 10 | 0 | -10 | -5 | F | F | F |
| 7 | 0 | 10 | -10 | 0 | -10 | F | T | F |
| 8 | 0 | 0 | 0 | 30 | 15 | T | T | F |
| 9 | 30 | 0 | 30 | -30 | 15 | T | F | T |
| 10 | 0 | 30 | -30 | NA | -15 | F | NA | NA |

This method is faster than the loop operation since pandas is based upon the numpy framework, which is insanely fast. This allows the pipeline to quickly call peaks.
