## Supplementary figures and images for "sRNAfrag: A pipeline and suite of tools to analyze fragmentation in small RNA sequencing data"

### Additional Figure 1

#### Deviation from True Loci

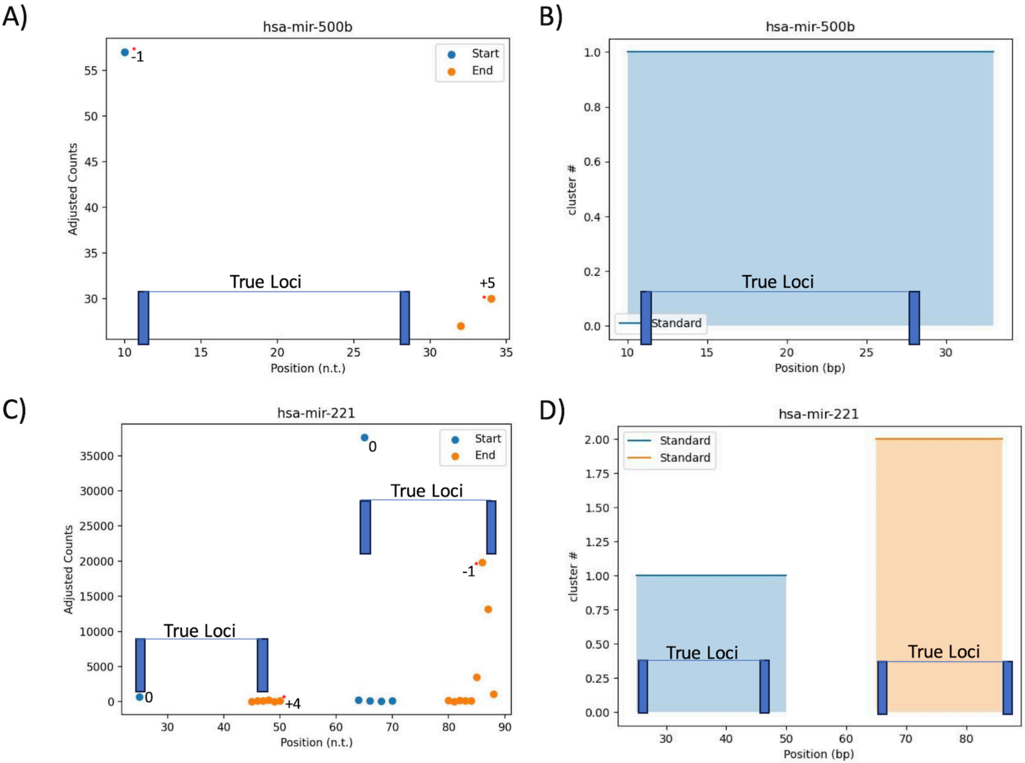

### Additional Figure 2

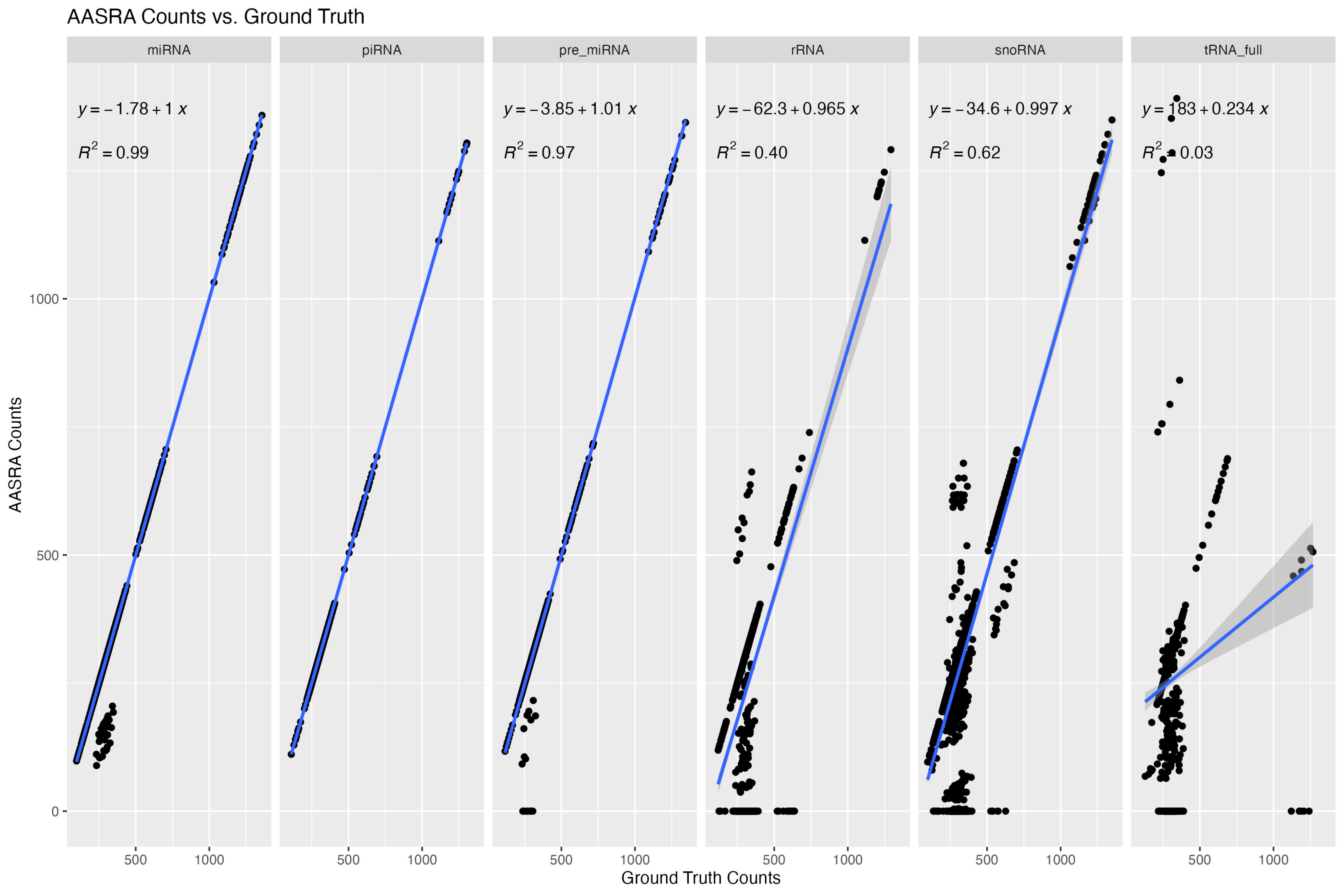
